# State-dependent protein-protein interactions mediating 4-1BB CAR Signaling

**DOI:** 10.1101/2022.06.07.495180

**Authors:** Samuel A. Ritmeester-Loy, Jonathan D Lautz, Yue Zhang-Wong, Joshua Gustafson, Ashley Wilson, Chenwei Lin, Philip R. Gafken, Michael C Jensen, Rimas Orentas, Stephen E.P. Smith

## Abstract

Cells rely on activity-dependent protein-protein interactions to convey biological signals, but the state-dependent interactome is notoriously cell-specific and undercharacterized^1^. In the case of chimeric antigen receptor (CAR) T cells containing a 4-1BB costimulatory domain, receptor engagement is thought to trigger the formation of protein complexes similar to those triggered by T cell receptor (TCR)-mediated signaling, but the number and type of protein-interaction-mediating binding domains differ between CARs and TCRs. Here, we performed co-immunoprecipitation mass spectrometry of a 2^nd^ generation CD19-directed 4-1BB:zeta CAR (referred to as bbζCAR) and identified 67 proteins that increased their co-association after target engagement. We compared activity-induced TCR and CAR signalosomes using quantitative multiplex co-immunoprecipitation and showed that bbζCAR engagement leads to activation of two modules of protein interactions, one similar to TCR signaling that is more weakly engaged in bbζCAR vs. TCR, and one composed of TRAF signaling complexes that is not engaged by the TCR. Batch-to-batch and inter-individual variations in IL2 production correlated with differences in the magnitude of protein network activation. Future CAR T cell manufacturing protocols could measure, and eventually control, biological variation by monitoring these signalosome activation markers.

**One Sentence Summary:** We define a network of protein interactions engaged by chimeric antigen receptors following target binding, and show that the magnitude of network activation correlates with IL-2 secretion, a proxy measure for CAR T cell function.

## Introduction

Chimeric antigen receptors (CARs) use a single-chain variable antibody fragment (scFv) specific to tumor antigens to instruct T cell activation via an engineered construct containing CD3ζ and costimulatory domains^2,3^. Four CARs targeting CD19, using either 4-1BB (axicabtagene ciloleucel^4^ and lisocabtagene maraleucel^5^) or CD28 (tisagenlecleucel^6^ and brexucabtagene autoleucel^7^) costimulatory domains are FDA-approved for B cell lymphomas, and clinical trials have reported up to 94% remission rates following therapy^8^. Despite these striking successes, many CAR design challenges remain. CARs are not very effective against solid tumors, likely due to the immunosuppressive tumor microenvironment, which prevents their more widespread use in oncology^9^. The selection of appropriate tumor antigens is critical to prevent on-target, off-tumor side effects, and tumor heterogeneity or down-regulation of antigens may lead to tumor escape and relapse. Even when a compatible tumor antigen is identified, as is the case with CD19 CARs, moderate-to-severe side effects such as cytokine release syndrome or neurotoxicity affect a significant portion of patients, which prevents the use of CAR T therapy as a first-line treatment^10^. Since the CAR is a synthetic receptor, it should be possible to bio-engineer our way around some of these issues. However, an engineering approach requires a solid understanding of the protein complexes that mediate downstream signal transduction and constitute the ‘programming language’ of the cell^11^.

The CAR is thought to instruct T cell activation by engaging protein-protein interactions similar to those engaged by the native T cell receptor/CD3 complex (TCR) in response to peptide-MHC stimulation^2^. In the TCR system, engagement of peptide/MHC leads to LCK- and FYN-dependent phosphorylation of CD3 ITAM motifs-one in each CD3δ, ε, and γ, and three in CD3ζ-which recruit SH2- and SH3-domain containing proteins such as ZAP70 in a phosphorylation-dependent manner^12^. In addition, accessory binding sites on the TCR, such as the NCK binding site in the proline rich region of CD3ε^13^ contribute additional complexity to the signalosome, rendering it sensitive to small changes in both peptide-MHC binding kinetics as well as the costimulatory environment. The overall strength of signalosome activation appears to dictate the cellular response^14^. By contrast, the CAR contains only a single CD3ζ intracellular domain, which allows the CAR to bind to ZAP70 following scFv engagement, facilitating its phosphorylation and activation^15^. However, mass spectrometry studies comparing the phospho-proteome following TCR vs. CAR stimulation have found that key signaling adapters downstream of ZAP70, including LAT and SLP76, are phosphorylated to a much lower degree, or not at all, following CAR vs. TCR engagement, suggesting less efficient signalosome formation^16,17^. In addition, the bbζCAR has a signaling motif not present in the TCR, derived from a TNF-receptor family protein (TNFRSF9 or 4-1BB) that trimerizes in response to ligand binding to activate downstream ERK and NFκB signaling via a mechanism involving the oligimerization of TNF Receptor Associated Factors (TRAFs) (reviewed in ^18^). TRAFs are required for 4-1BB enhancement of bbζCAR function^19^, through canonical or non-canonical NFκB pathways^20^. However, the specific activity-dependent protein complexes recruited to the CAR following ligand binding, and their batch- and patient-specific variability, have not been thoroughly described.

We immunoprecipitated the bbζCAR and used mass spectrometry to compare co-associated proteins before and after CAR stimulation. We identified 67 proteins that increased their co-association with the bbζCAR following CD19 stimulation, including ZAP70 and a large TRAF signaling complex. We incorporated the TRAF signalosome into a quantitative multiplex co-immunoprecipitation (QMI) platform previously used to study dynamic protein-protein interactions downstream of T cell receptor activation^14,21^, and compared signalosome formation following activation of TCR and bbζCAR. We found a core network of interacting proteins that changes its pattern of co-association following CAR ligation, which varied in composition and intensity of activation between batches, CAR scFv targets, and individual donors. Our results support a model in which bbζCAR recruits ZAP70, which induces the formation of a LAT/SLP76 signalosome that is quantitatively weaker than that induced by TCR stimulation, while simultaneously engaging TRAF-mediated complexes that include BIRC2/3 and TAK1. Batch- and patient-specific differences in the intensity of signalosome activation correlate with IL-2 production, which suggests that the expression or assembly of key signalosome components may contribute to variations in CAR performance.

## Results

### Mass spectrometry identification of signaling-induced protein complexes

To identify a bbζCAR-specific signalosome, we immunoprecipitated the protein complexes bound to the bbζCAR following ligand binding and compared them to those immunoprecipitated in the basal (unstimulated) state (Fig 1A). For maximal clinical relevance, we used CD4+ and CD8+ T cells from four healthy donors lentivirally transduced in a GMP facility with a clinical bbζCAR containing an anti-CD19 scFV, an IgG4 hinge, a CD28 transmembrane region, a 4-1BB costimulatory domain, the complete intracellular sequence of CD3ζ, and a T2A self-cleaving peptide that co-expresses a truncated EGFR marker and leaves a small “2A scar” on the C-terminus of the mature CAR. This CAR has undergone extensive clinical trials at our local institution (NCT02028455)^8^, and is identical to the FDA-approved lisocabtagene maraleucel^5^ product. Western blotting demonstrated successful IP of the CAR, and co-immunoprecipitation of ZAP70 in the stimulated condition only indicated successful detection of activity-dependent binding (Fig 1B).

**Figure 1:**
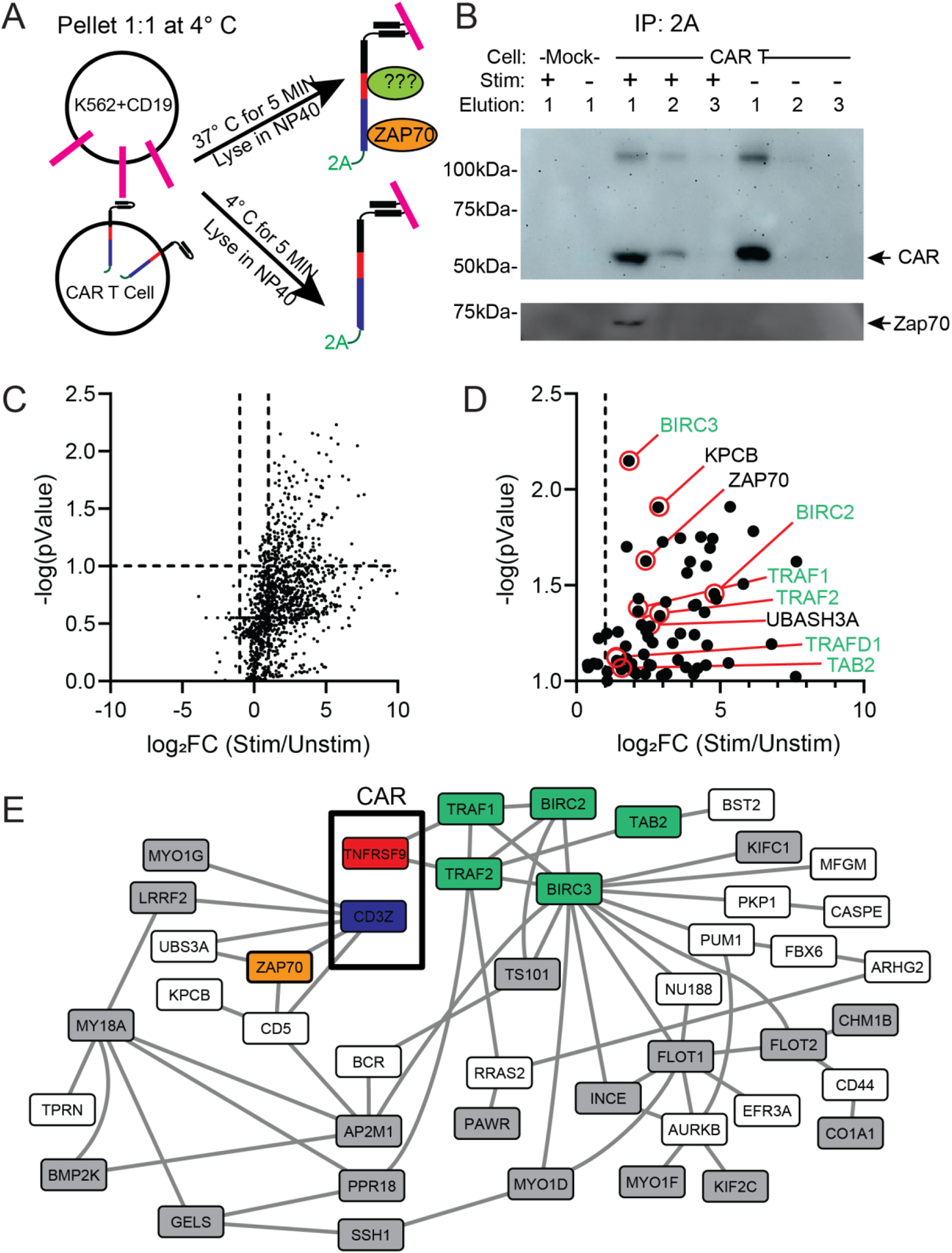
Activity-dependent CAR interactome identified by IP-mass spectrometry. A) Experimental design. B) IP-Western blot demonstrating successful co-IP and complete elution of CAR, as well as co-association of ZAP-70 exclusively in the stimulated condition. Representative of at least N=4 separate blots. C) Volcano plot showing all proteins detected by LC/MS/MS. D) Upper-right quadrant of volcano plot after removing CRAPome proteins, with selected proteins highlighted. E) Known protein-protein interactions among proteins identified in D. Green coloring indicates TRAF signaling complex, and grey indicates cytoskeletal-related proteins. Note that detected proteins with no database interactions are not shown (e.g. TRAFD1).

Mass spectrometry identified a total of 2,280 proteins in four datasets (N = 22 total samples derived from four individuals; 3 CD8 / 4 CD4 CAR T cells; 4 mock-transduced T cells; each ‘stimulated or ‘not stimulated). To identify activity-dependent interactions, we compared proteins in the stimulated vs. unstimulated condition for combined CD4 and CD8 cells using a mixed linear model, and stringently removed “noise” proteins that appeared in more than 10% of human affinity purification experiments in the CRAPome database^22^. Using cutoffs of false discovery rate (FDR)<0.1 and a log_2_ fold change (log_2_FC)≥1, we identified 67 proteins that were significantly increased after stimulation (Fig 1C-D, Table S1). Importantly, ZAP70 was among the identified proteins (FC=5.3, t=−2.77). The most significantly upregulated protein was BIRC3 (FC=3.5, t = −3.58), an E3 ubiquitin ligase involved in TNF receptor family signaling. Ligand-induced trimerization of TNF-family receptors (such as 4-1BB) leads to recruitment of BIRC3, as well as its homologue BIRC2, which ubiquitinate TRAF family proteins and recruit additional signaling molecules including TAB1/2 and TAK1 (aka MAP3K7), leading to initiation of MAPK and NFKB signaling^18^. Six members of this signalosome were among our 67 identified ligand-dependent interactors: TRAF1 (FC= 4.4, t=−2.40), TRAF2 (FC= 7.4, t=−2.36), TRAFD1 (FC= 2.8, t=−2.01), BIRC2 (FC= 27.8, t=−2.52), BIRC3 (FC=3.5, t = −3.58), and TAB2 (FC= 2.6, t=−2.01). Other notable proteins included PKCβ (KPCB, FC=7.1, T=−3.19), a kinase that mediates the activation of NFκB by phosphorylating CARD11/CARMA and forming a complex that includes TAK1^23^; UBASH3A (FC=5.9, T=−2.28), which regulates ubiquitination and degradation of the TCR signalosome by inhibiting CBL-B^24^; CD5 (FC=2.1, T=−2.23), a negative regulator of TCR signaling that binds CBL-B and UBASH3A, dampens NFκB signaling, and promotes effector/memory phenotypes^25,26^; and CD44 (FC=8.0, T=−2.92), a marker of memory/effector T cells that binds LCK and may potentiate TCR responses by recruiting LCK to signaling sites^27^.

We next used the PICKLE database^28^ (Fig 1E) to visualize previously reported protein-protein interactions among our identified proteins, as well as the STRING database^29^ to highlight functional (including non-physical) associations (Fig S1). We included known TNFRSF9 and CD3Z interactions to represent the bbζCAR. The PICKLE network showed a high degree of connectivity, centered around TRAF signaling; BIRC3 (11 edges) and TRAF2 (9 edges) were the most connected nodes. ZAP70 had known interactions with UBASH3A, CD5, and DOCK8, a guanine nucleotide exchange factor (GEF) that activates CDC42, is required for CD4+ T cell migration in response to chemokine^30^, and regulates NK cell cytotoxicity^31^. The most enriched KEGG pathway was “NFkB signaling pathway” (p = 3.43 × 10^−5^), followed by “apoptosis” (p = 0.001) and TNF signaling pathway (p = 0.0048) (Fig S1). T cell receptor signaling pathways were not significantly enriched; in fact, it was somewhat surprising that we did not identify more T cell receptor signalosome-associated proteins: LCK (FC= 2, t=−1.15, not significant (NS)), SLP76 (not detected), LAT (not detected), GRAP2 (FC = 9.2, t=1.19 NS), CD28 (FC = 1.03, t= −1.15 NS) and other major T cell signaling proteins did not reach statistical significance. We also identified a large number of actin binding proteins (grey in Fig 1E), including proteins related to Ras/Rac/CDC42 signaling, suggesting that the activated CAR may engage actin cytoskeletal rearrangement, and many of the top gene ontology (GO) processes involved cytoskeleton organization (p = 5.45×10^−10^) and actin filament-based processes (p = 1.23×10^−9^) (Fig S1). While actin is a frequent contaminant in mass spectrometry experiments, these associations were still detected after removal of cytoskeletal elements present in the CRAPome database. Overall, the interactors identified highlight extensive cytoskeletal rearrangements, ZAP70-mediated interactions not traditionally associated with TCR signaling, and the activation of TRAF-mediated signaling complexes.

### TCR stimulation measured in mock and bbζCAR T cells by quantitative multiplex co-immunoprecipitation

Quantitative multiplex co-immunoprecipitation (QMI) uses antibody-coupled flow cytometry beads to immunoprecipitate protein complexes, and fluorophore-coupled probe antibodies directed to different protein targets to monitor acute changes in protein co-association using a flow cytometer^21,32^. QMI measures Proteins in Shared ComplexES (PiSCES), so to maximize protein complex detection, so we added several of the most connected nodes in Fig 1E to our previously-described QMI panel, which was built to measure the amount of co-association among critical TCR signalosome members^14,21^: CAR (anti-2A scar, IP only), TRAF1, TRAF2 and BIRC3. We also included three proteins that did not reach statistical significance in the mass spec experiments, but are known to be important to the TRAF and NFκB signalosome in lymphocytes: TAK1 (MAP3K7, FC=597, t=−1.09 NS), which is recruited by TAB1/2 and signals to IKKb^18^ and was recently shown to be critical to IFNγ production in CD8+ T cells in a large CRISPR screen^33^; TNIK (found in only 1 co-IP experiment) which binds TRAFs, may recruit NCK to link TRAFs with TCR signaling mechanisms^13^, and is critical for CD8+ T cell memory formation^34^; and SHARPIN (FC=13.6, T=−1.21), a component of the linear ubiquitination complex (LUBAC) that promotes activation of the IKK complex downstream of TNF family receptors^35,36^, ubiquitinates CARMA-BCL10-MALT complex^37^, and may bind directly to TRAF1^38^. As previously described^21,32,39^, we identified two antibodies that could simultaneously bind each target in its native state on flow cytometry beads, and validated target specificity using cell lysates lacking the target (Fig S2 and Table S2). Detergent optimization is critical to co-immunoprecipitation experiments, but with multiplex co-IPs, one must select the best compromise detergent for the 100s of potential binary PiSCES measurements^40^. We compared Digitonin (DIG), a detergent used in studies of TCR signaling due to its ability to maintain native TCR/CD3 complexes, with NP-40, a detergent used in studies of CAR signaling that can better solubilize the CAR compared to DIG. We found a strong effect of detergent on detected protein networks; while many PiSCES were detected in both detergents, others were better detected in NP-40 (e.g. CAR_ZAP70) or in DIG (e.g. SLP76_LAT) (Fig S3). We therefore began our initial characterizations of CAR signaling using both detergents, NP-40 in Fig 3, and DIG in Figs 2/4/5.

**Figure 2).**
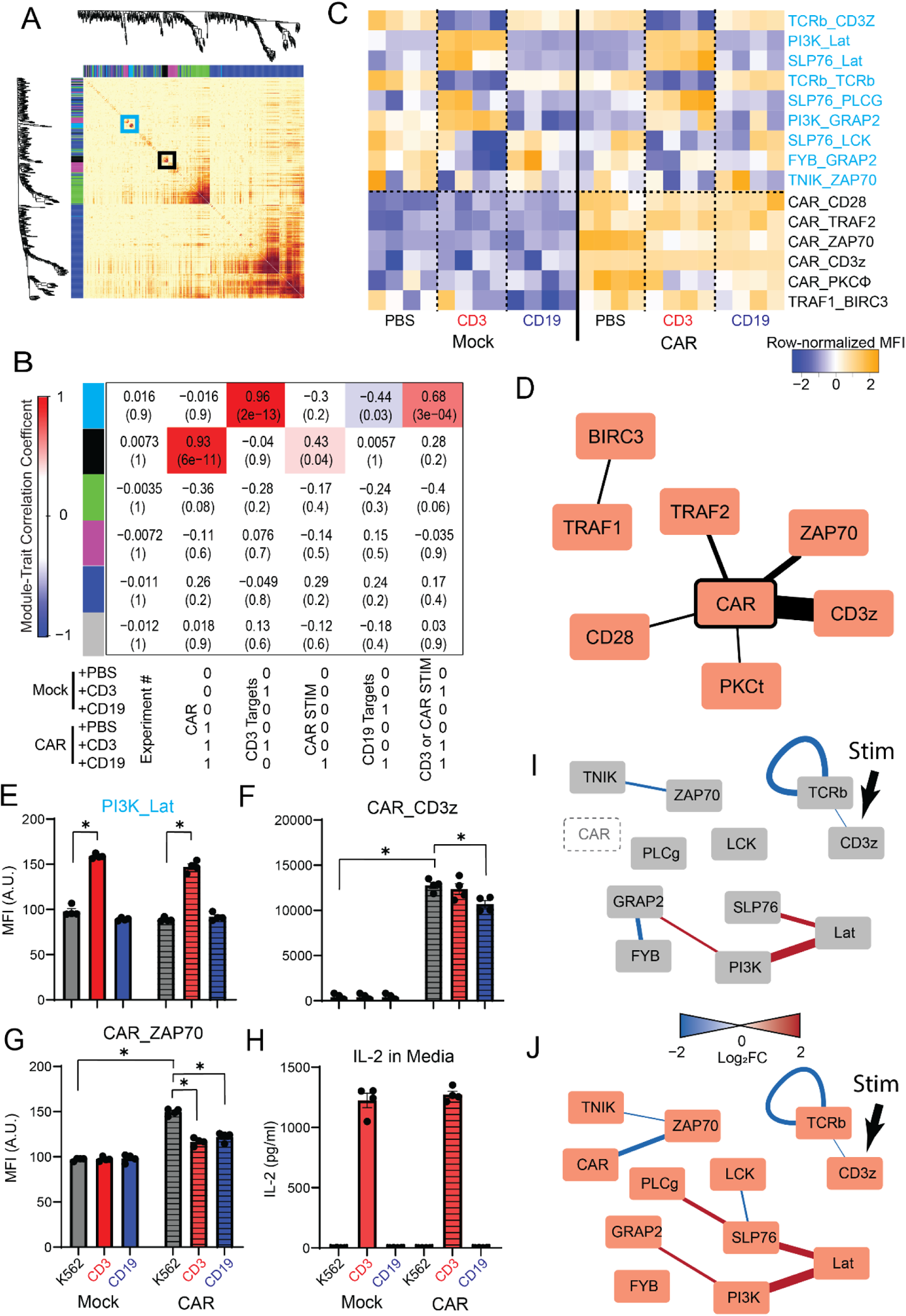
TCR stimulation is intact in CAR T cells. bbζCAR or mock-transduced T cells were stimulated with platebound anti-CD3 (OKT3) or CD19 for 5 minutes. A) Correlation network analysis clusters all detected PiSCES by their correlated behavior across N=24 individual measurements, and identifies color-coded modules of correlated interactions. B) A module-trait correlation table reports the correlation coefficient and p-value (in parenthesis) between the eigenvector of each color-coded CNA module (colored boxes) and a binary description of experimental variables, below table. Module-trait correlations greater than 0.4 are highlighted with red (positive correlation) or blue (negative correlation). C) Row-normalized heatmap of ANCՈCNA-significant proteins in shared complexes (PiSCES) in the turquoise or black modules. D) Proteins in the black module co-associate with the unstimulated CAR. Edge thickness indicates relative intensity of PiSCES detection. E) Median fluorescent intensity (MFI) of PI3K_LAT, representative of the turquoise module, which increases following anti-CD3 stimulation. * p<0.05 by ANC. F) MFI of CAR_CD3, representative of the Black module, which increased with CAR expression and decreased with anti-CD19 stimulation. * p<0.05 by ANC. G) MFI of CAR_ZAP70 increased with CAR expression, but decreased with anti-CD3 or CD19 stimulation. * p<0.05 by ANC. H) IL2 secretion into media after overnight incubation on anti-CD3 but not CD19-coated plates. I,J) PiSCES that changed with anti-CD3 stimulation in I) mock-transduced or J) CAR T cells. Edges represent a change in the amount of interaction between the connected nodes; edge color and width indicate direction and magnitude of the change.

In order to observe dynamic protein networks without the confounding factor of target cells being present, we stimulated primary human CAR T or mock cells for 5 minutes with platebound anti-CD3/CD28 or CD19, and lysed in digitonin for consistency with prior TCR signaling studies^14,21^. Correlation network analysis (CNA)^41^ identified 6 modules of PiSCES whose behavior correlated across experiments (Fig 2A), including a ‘turquoise’ module that correlated with CD3 stimulation (correlation coefficient (CC)=0.96, p=2×10^−13^), and a ‘black’ module that correlated with both CAR presence (CC=0.93, p=6×10^−11^) and to a lesser degree, CAR stimulation (CC=0.43, p=0.04) (Fig 2B). To filter out noise and ensure only the most robustly changed PiSCES were reported, a custom-built adaptive, non-parametric statistical test corrected for multiple comparisons (ANC, see ^21^) was used to compare PiSCES in different conditions, and only PiSCES that were both ANC-significant, and significantly correlated with a CNA-significant module (ANCՈCNA PiSCES) are reported^14,21,39^.

Anti-CD3 stimulation led to alterations in 9 turquoise-module PiSCES in both cell types (Fig 2C). The complex with the highest module membership (MM) in the turquoise module (i.e. most strongly correlated to the eigenvector of the module), TCR_CD3, was markedly reduced following platebound stimulation, while PI3K_LAT (Fig 2E) and PiSCES containing GRAP2 and SLP76 increased, consistent with formation of a TCR signalosome^21^. In fact, PI3K_LAT was the most upregulated PiSCES following peptide/MHC stimulation of naïve, double-positive mouse thymocytes^14^. Importantly, the response to anti-CD3 was similar in CAR-expressing and mock-transduced T cells (Fig 2I, J), indicating that CAR expression did not interfere with TCR-mediated signalosome formation.

ANCՈCNA PiSCES in the black module (Fig 2D), were increased with CAR expression, indicating formation of these PiSCES in unstimulated CAR T cells. Complexes including CAR_TRAF2, CAR_PKCΦ, and CAR_ZAP70 were apparent. In response to plate-bound CD19, only two PiSCES were altered: CAR_CD3ζ (Fig 2F) and CAR_ZAP70 (Fig 2G). It is important to note that the CD3ζ probe antibody, 6B10.2, reacts to a portion of CD3ζ that is included in the CAR construct^42^, so this PiSCES may measure immunoprecipitation and/or multimerization of the CAR itself, or an interaction between the CAR and a native CD3ζ component such as the CD3ζ “p21” protein that co-associates with 2^nd^-generation CARs^43^. CAR_ZAP70 decreased with plate-bound CD19 exposure, which is not consistent with CAR stimulation because CD19 stimulation increases this co-association (e.g. Fig 1B). Moreover, CAR T cells produced no IL-2 following overnight incubation with plate-bound CD19, but did following anti-CD3 (Fig 2H), consistent with a failure of plate-bound CD19 to engage CAR signaling complexes. Overall, these data revealed that CD3 stimulation induced protein complexes containing LAT, PI3K, SLP76, GRAP2 and others in both Mock T cells and CAR T cells (Fig 2I,J), and further demonstrated PiSCES that formed following CAR expression (Fig 2 D). However, plate-bound CD19 stimulation was unproductive.

### Activity-induced changes in TCR and CAR signaling networks

We next stimulated bbζCAR T cells with anti-CD3- or CD19-expressing K562 target cells that were briefly fixed with glutaraldehyde to prevent the target cell’s protein content from interfering with QMI detection. A similar strategy was used previously for stimulation of mouse and human TCRs with peptide-MHC^14,21^. Proteins were solubilized in NP-40 to ensure complete solubilization of the CAR. CNA identified a turquoise module that correlated with CD3 stimulation (CC=0.92, p=1×10^−10^), and also correlated to a lesser degree with CD3 and CD19 stimulation (CC=0.74, p=4 × 10^−5^) (Fig 3A). ANCՈCNA identified 17 PiSCES that increased following CD3 stimulation in both CAR and mock T cells (Fig 3B). Increased detection of the PiSCES with the highest MM, LCK_ZAP70 (Fig 3C), as well as CD3ζ_LCK, indicated activation of the CD3ζ-LCK-ZAP70 cascade typical of TCR signaling, and increased PI3K_LAT (Fig 3D) and SLP76_LAT indicated transduction of the ZAP70 activation to a LAT-mediated signaling complex. Interestingly, TRAF2_TRAF2, TAK1_TAK1 and TNIK_TNIK levels were also increased. Since proteins are unlikely to be synthesized de novo in 5 minutes, this increase in QMI detection may indicate changes in self-association or in the accessibility of antibody epitopes, and suggests conformational changes in TRAF signaling complexes downstream of TCR/CD3 stimulation. ANCՈCNA node-edge diagrams for CAR and mock T cells were similar, although not identical due to biological noise and the stringency of the statistical analysis, and again demonstrate that CAR expression did not alter TCR-mediated signaling (Fig 3G, H).

**Figure 3:**
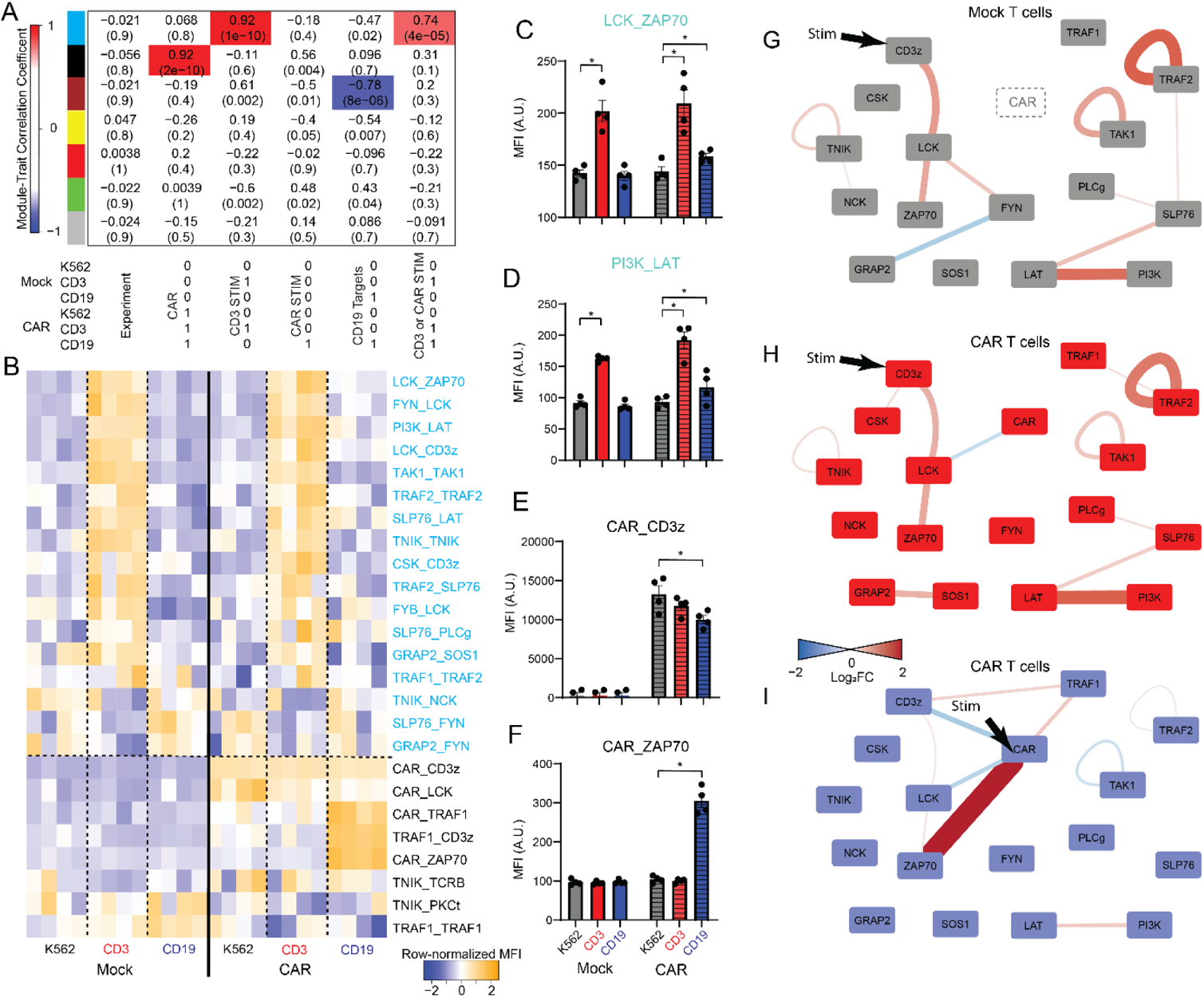
bbζCAR activates TCR- and TRAF-associated signaling networks. bbζCAR or mock-transduced T cells were stimulated with fixed K562 targets expressing nothing (parental), anti-CD3 (OKT3) or CD19 for 5 minutes. A) A module-trait correlation table reports the correlation coefficient and p-value (in parenthesis) between the eigenvector of each color-coded CNA module (colored boxes) and a binary description of experimental variables, below table. Module-trait correlations greater than 0.6 are highlighted with red (positive correlation) or blue (negative correlation). B) Row-normalized heatmap of ANCՈCNA-significant proteins in shared complexes (PiSCES) in the turquoise or black modules. C-F) MFI of (C) LCK_ZAP70 and (D) PI3K_LAT representative of the Turquoise module, which increases following anti-CD3 stimulation and increases to a lesser degree following CD19 stimulation. * p<0.05 by ANC. MFI of (E) CAR_CD3 and (F) CAR_ZAP70, both in the Black module. G-I) PiSCES that changed with anti-CD3 stimulation in G) mock-transduced or H) CAR t cells, or that changed with I) CD19 stimulation in CAR T cells. Edges represent a change in the amount of interaction between the connected nodes; edge color and width indicate direction and magnitude of the change.

Following CAR stimulation, three PiSCES in the turquoise module, LCK_CD3ζ, LCK_ZAP70 and PI3K_LAT, were also significantly increased, demonstrating partial, although considerably weaker, activation of ZAP70-to-LAT signaling downstream of CAR ligation (Fig 2, C,D). In addition, a second module, Black, correlated with CAR expression (CC=0.92, p=2×10^−10^), and to a lesser extent, CAR stimulation (CC=0.56, p=0.004). The PiSCES with the highest MM, CAR_CD3ζ, was strongly detected in unstimulated CARs, and was significantly reduced following stimulation (Fig 3E). Similarly, CAR_LCK was significantly reduced. CAR-ZAP70 (Fig 3F), CAR_TRAF1, and TRAF1_CD3ζ increased following CAR ligation, and the abundance of TRAF2_TRAF2 and TAK1_TAK1 decreased, indicative of changes to their multimerization and/or binding partners/antibody accessibility. Overall, these data indicate the rapid formation of a CAR-ZAP70 signaling complex, which activates TCR-signalosome components to a lesser degree than TCR-mediated signaling, and simultaneous engagement of a TRAF signaling complex unique to the CAR.

### Paradoxically reduced signaling complexes detected by QMI

We expected PiSCES such as CAR_LCK or CAR_TRAF2 to be increased, not decreased, following CD19 ligation, since signalosome activation is thought to involve assembly of protein complexes. However, QMI queries protein complexes in the native state, so the apparent reduction in PiSCES abundance may reflect a true reduction in the measured co-association, or may reflect an inability of a probe antibody to bind a target decorated with activity-dependent interactions or ubiquitin chains. It may also reflect the CAR irreversibly binding to fixed target cells and being excluded from the soluble lysate. In order to discount this latter possibility, we repeated the fixed-target experiment (Fig 3) with unfixed targets, and obtained similar results (Fig S4). In this experiment, the CAR showed increased apparent co-association with TRAF1 and ZAP70, but decreased co-association with CD3ζ, LCK and TRAF2 (Fig S4). PI3K_LAT was increased, while other CD3-responsive PiSCES were not significantly changed. We also performed co-IP western blots of the CAR before and after stimulation with fixed targets, and found equal amounts of CAR in the immunoprecipitate (Fig S5), suggesting a conformation explanation for the apparent reduction in CAR_CD3ζ. When western blotting was able to detect co-IP’d TRAF2 (which did not occur in all experiments), we found an increase in the amount of TRAF2 co-associated with the CAR (Fig S5), consistent with mass spectrometry results (Fig 1) and previous reports (e.g. Fig 5D in ref ^44^). These data reveal a consistent pattern of changes in CAR co-associations using fixed or unfixed targets and discount the hypothesis that the activated CAR is depleted from the lysate by binding to fixed targets. Our data support a model that, following CD19 engagement, recruited protein complexes or ubiquitination events sterically hinder our probe antibodies, resulting in paradoxically reduced complex detection by QMI.

### Consistency of bbζCAR signaling complexes

Differences in the behavior of CARs targeted to different antigens are well-established, and can be due to differences in tonic activation, the size or abundance of the protein target, or small differences in CAR design^45^. In addition, batch-to-batch variation inherent to the manufacturing process may contribute to variable clinical responses^46^. In order to quantify similarities and differences in CAR signaling due to batch and target, we generated four batches of CAR T cells from a single donor, two with an anti-CD19 scFv, and two with an anti-EGFR“806” scFv^47^. The CD19CAR and EGFR806CAR are identical in structure except for the specificity-determining scFv.

Hierarchical clustering of the data matrices showed samples clustering first by CAR stimulation, then by CAR type (anti-CD19 vs anti-EGFR), and finally by batch (Fig 4A). CNA identified two modules that correlated with CAR stimulation: turquoise (CC = 0.62, p=4×10^−4^) and black (CC= −0.64, p=1×10^−4^) (Fig 4B). The ANCՈCNA PiSCES with the highest module membership in turquoise was PI3K_LAT (Fig 4D), followed by SLP76_LAT, indicating consistent formation of the LAT signalosome following CAR engagement (Fig 4 B,C). CAR_ZAP70 (increased, Fig 4D), CAR_PKCΦ (decreased, Fig 4E), and CAR_SHP2 were also members of the turquoise module, suggesting that these PiSCES contribute to the formation of a ‘TCR-like signalosome’ downstream of CAR ligation. The black module PiSCES with the highest MM were TRAF-related, including BIRC3_BIRC3 (Fig 4F), TRAF1_BIRC3, and CAR_TRAF2 (Fig 4G). Upon bbζCAR engagement, the apparent abundance of these protein complexes was reduced, indicative of a rearrangement of TRAF signaling components that contribute to the orchestration of bbζCAR signaling, as discussed above. These data define a core set of protein complexes, organized into two modules, that mediate bbζCAR signaling (Fig 4J).

**Figure 4:**
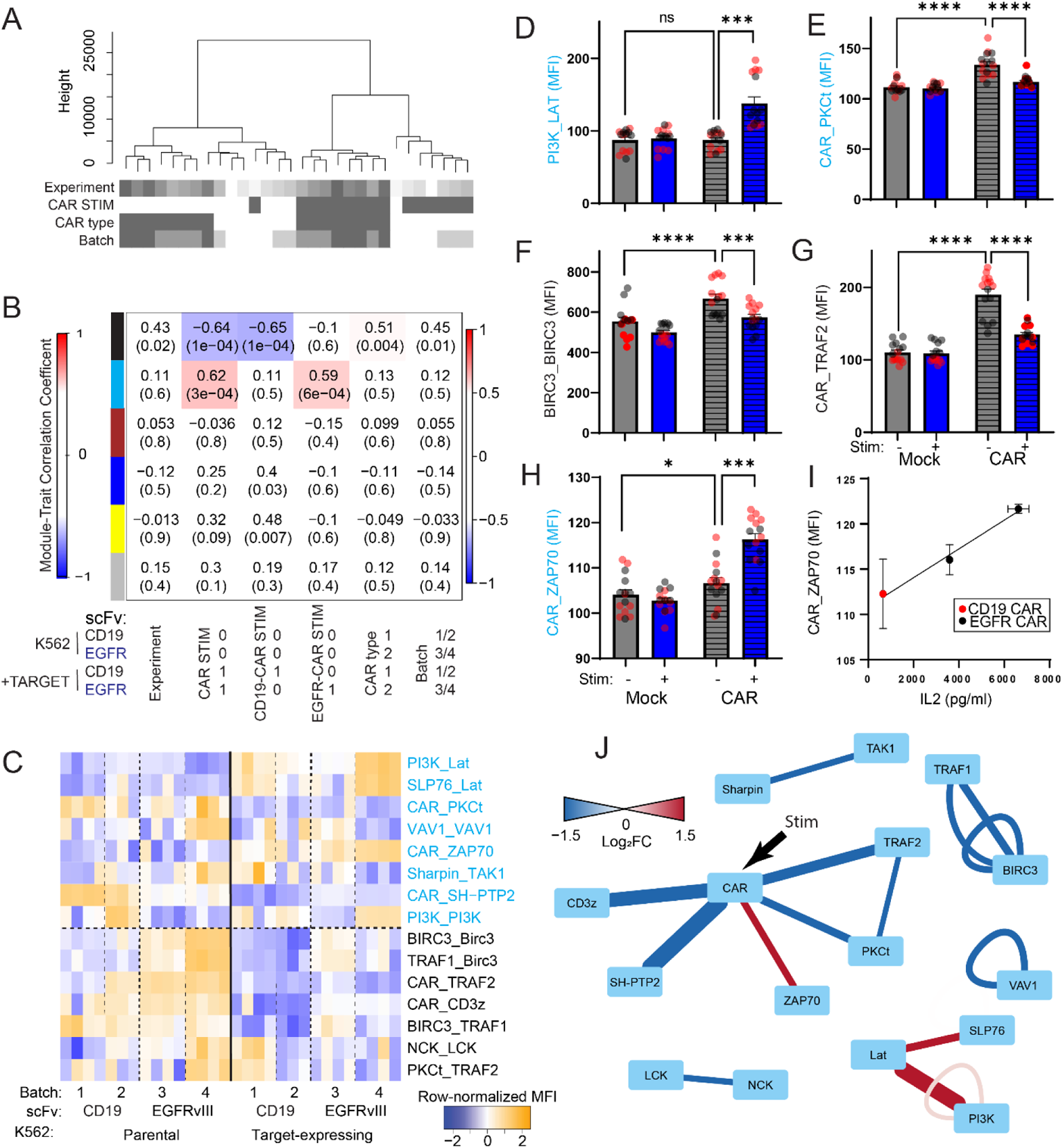
Consistent bbζCAR signaling despite batch- and target-specific variability. Four batches of bbζCAR, with two different scFv targets, were manufactured from a single donor and lysed in DIG. A) Hierarchical clustering of all detected PiSCES shows clustering by stimulation, then by CAR type, and batch. B) A module-trait correlation table reports the correlation coefficient and p-value (in parenthesis) between the eigenvector of each color-coded CNA module (colored boxes) and a binary description of experimental variables, below table. Module-trait correlations greater than 0.5 are highlighted with red (positive correlation) or blue (negative correlation). C) Row-normalized heatmap of ANCՈCNA-significant proteins in shared complexes (PiSCES) in the turquoise or black modules. D-H) MFI of (D) PI3K_LAT and (E) CAR_PKCΦ, members of the Turquoise module, and (F) BIRC3_BIRC3 and (G) CAR_TRAF2, members of the black modeule, and H) CAR_ZAP70, following CD19 stimulation of mock-transduced or bbζCAR T cells. * p<0.05 by one-way ANOVA followed by Sidak’s multiple comparison test. I) Correlation between the MFI of CAR_ZAP70 and IL-2 in the media 18 hours later in CD19-stimulated in CAR T cells. J) Node-edge diagram showing PiSCES significantly changed (by ANCՈCNA) following bbζCAR target engagement. Edges represent a change in the amount of interaction between the connected nodes; edge color and width indicate direction and magnitude of the change. N = 60 biological replicates, 15 per condition. Only CAR+ cells (N=15 unstim, 15 stim) were included in the analyses represented in A, B, C and J.

While we were able to detect PiSCES that changed in all batches, there were also clear differences between the four batches of CAR T cells (e.g. PI3K_LAT and CAR_ZAP70 (Fig 4C)) due to first target type, then manufacturing run. Uncontrollable differences between CAR production batches may contribute to clinical CAR performance, one indicator of which is IL-2 secretion following overnight incubation with targets. Remarkable, IL-2 secretion correlated with the average MFI of CAR_ZAP70 after CD19 exposure (Fig 4I), with the batches showing the strongest signalosome activation also showing the strongest IL-2 production. These data suggest batch-specific signalosome activation may predict batch-specific CAR performance.

### Donor-dependent differences in CAR signaling predict IL-2 production

Until this point, all experiments were performed on CAR T cells manufactured from a single donor to eliminate genotype-dependent variation. To investigate individual variation, we made four batches of anti-CD19 bbζCAR from four donors and stimulated with fixed K562 target cells expressing anti-CD3 or CD19. CNA identified two modules, turquoise and black, which correlated with both anti-CD3 stimulation (MM = −0.91, p = 9 × 10^−9^) and CD3 or CD19 stimulation (MM = −0.73, p = 6 × 10^−5^), or with CAR expression (MM = 0.82, p = 7 × 10^−7^), respectively (Fig 5A, B). The turquoise module contained PiSCES representing the TCR complex, including TCR_CD3ζ (Fig 5C), which were not detected in experiments lysed in NP40 (e.g. Fig 3, see Fig S3), as well as PI3K_SLP76 and PI3K_LAT (Fig 5C), representing TCR signalosome formation. The black module contained CAR and TRAF-containing PiSCES including CAR_TRAF2 (Fig 5E), which reduced their apparent abundance following CAR stimulation. The intensity of CAR_ZAP70 (Fig 5F) and PI3K_LAT correlated with IL-2 secretion measured in overnight culture, again indicating signalosome formation may be predictive of performance. Node-edge diagrams showed similar signalosome formation in mock and CAR T cells downstream of CD3 stimulation (Fig 5G, H), which shared PI3K-LAT-SLP76 signalosome components with CAR stimulation (Fig 5I). These data demonstrate signalosome formation downstream of TCR and CAR which is qualitatively similar but quantitatively different among different donors. These intensity differences correlate with a functional readout of CAR activity.

**Figure 5:**
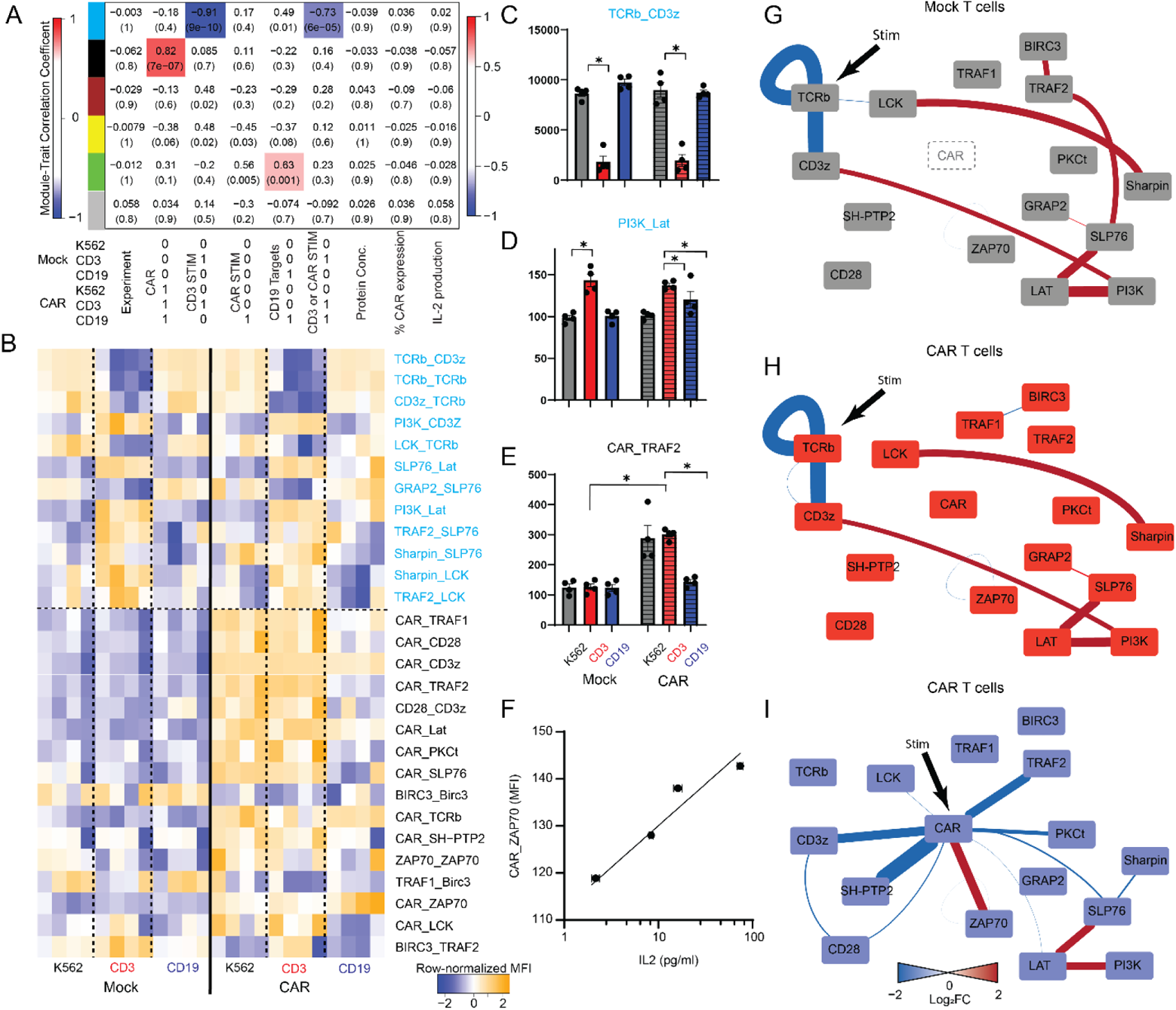
CAR signalosome dynamics are consistent among donors and correlate with IL2 secretion. Four batches of anti-CD19 bbζCAR, were produced from four donors and lysed in DIG. A) A module-trait correlation table reports the correlation coefficient and p-value (in parenthesis) between the eigenvector of each color-coded CNA module (colored boxes) and a binary description of experimental variables, below table. Module-trait correlations greater than 0.6 are highlighted with red (positive correlation) or blue (negative correlation). B) Row-normalized heatmap of ANCՈCNA-significant proteins in shared complexes (PiSCES) in the turquoise or black modules. Data are grouped by mock or CAR t cells indicated by the solid black line, then target type indicated by the dashed lines; the first column following each line represents matched conditions from donor 1, the second donor 2, etc. C-E) MFI of (C) TCR_CD3ζ and (D) PI3K_LAT, members of the Turquoise module, and (E) CAR_TRAF2, a member of the black modeule, following CD19 stimulation of mock-transduced or bbζCAR T cells. * p<0.05 by ANC. F) Correlation between the MFI of CAR_ZAP70 and IL-2 in the media 18 hours later in CD19-stimulated in CAR T cells. G-I) Node-edge diagram showing PiSCES significantly changed (by ANCՈCNA) following anti-CD3 stimulation in G) mock-transduced or H) CAR t cells, or I) CD19 stimulation in CAR T cells. Edges represent a change in the amount of interaction between the connected nodes; edge color and width indicate direction and magnitude of the change. N = 24 biological replicates, 4 per condition, derived from 4 different de-identified healthy donor.

## Discussion

We detected two modules of coordinated protein interactions that changed their pattern of co-associations following CAR activation: a TCR-like module, and a TRAF module (Fig 6). TCR activation is initiated by the kinase LCK, some of which is bound to a CXCP motif on coreceptors CD4 or CD8a^12^ and brought in physical proximity to the TCR when these coreceptors interact with MHC. Free, membrane-tethered LCK, not bound to coreceptor, may also play a role in basal TCR phosphorylation and activation^48^, as may LCK bound to the TCR through an interaction involving the CD3ε BRS motif^49^. The CAR does not engage CD4 or CD8, nor does it contain a CD3ε motif, so the origin of activating LCK is assumed to be the free LCK pool, which has higher kinase activity than the bound pool^48^. Irrespective of the source of LCK, the next step in both T cell and CAR activation is the recruitment of ZAP70 to LCK-phosphorylated ITAMs, and its subsequent phosphorylation by LCK, leading to ZAP70 activation. In the TCR system, LCK binds simultaneously to TCR-bound, activated ZAP70 and to the scaffolding adapter protein LAT, increasing the efficacy of the signaling cascade^50^.

**Figure 6:**
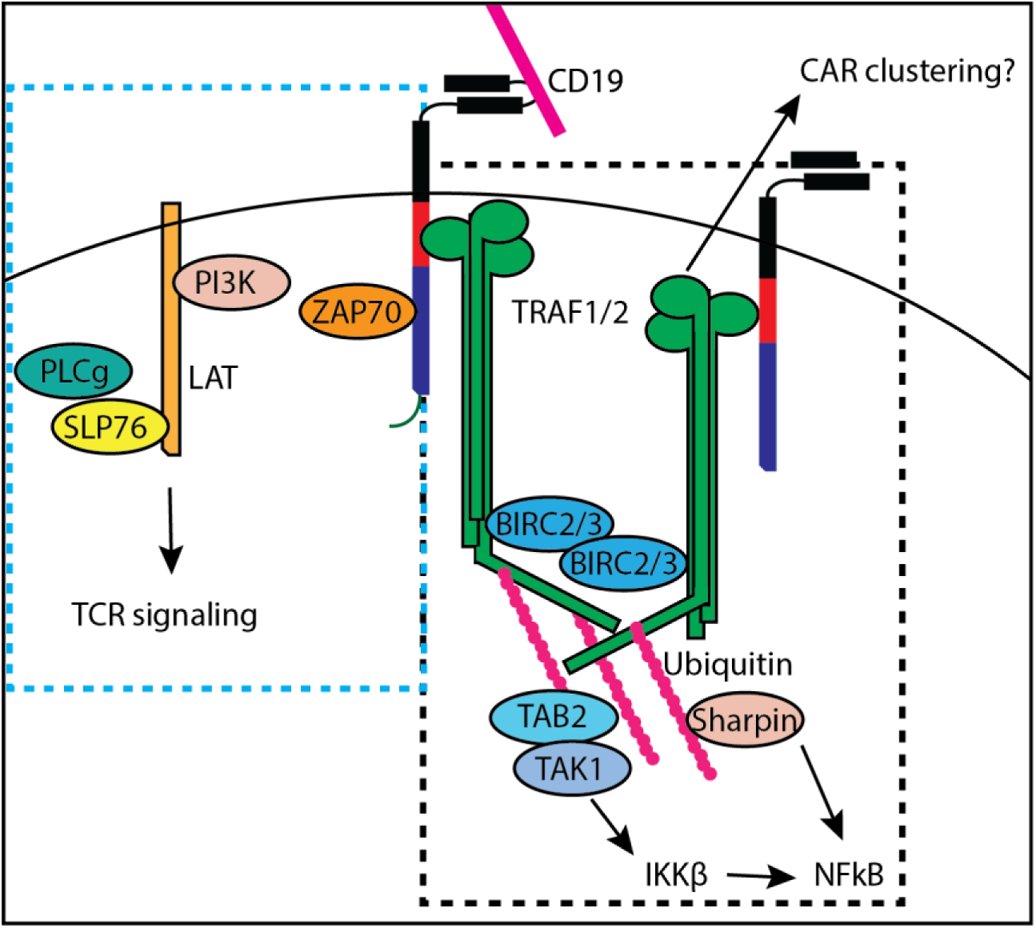
Proposed model of CAR signaling. Upon CD19 engagement, the bbζCAR engages two modules (dashed boxes), one composed of SLP76-LAT-PI3K signaling complexes that mimics TCR signaling, the other composed of TRAF signaling complexes that initiates NFkB signaling.

Phosphorylated LAT assembles a signalosome consisting of GRAP2, SLP76, PLCγ1, PI3K, and many other signaling molecules to collectively amplify TCR signals and initiate calcium signaling, MAPK activation, and actin polymerization^12^. Following OKT3 stimulation of the TCR, QMI detected increased co-association of CD3_LCK, LCK_ZAP70, LAT_SLP76, PI3K_LAT, SLP76_PLCg, and others, indicative of canonical TCR activation, in both CAR and mock T cells stimulated with OKT3. Following CAR stimulation, PI3K_LAT and SLP76_LAT was detected, but the magnitude of activation was lower compared to TCR stimulation, and the other PiSCES were not detected. Indeed, there were few other canonical TCR-associated proteins that associated with the CAR following stimulation, and “T cell signaling” was not an enriched GO or KEGG pathway. These data are consistent with prior studies on CAR T cells showing less efficient LCK engagement and ZAP70 mobility^15^ and less ZAP70-mediated phosphorylation of downstream adaptors such as PLCg and LAT^17^. The QMI assay can acutely measure this inefficient TCR-associated signalosome formation following CAR activation, offering a platform to monitor CAR performance and screen for improved CAR designs.

A second signaling module identified by mass spec and QMI encompasses TRAF signaling, consisting of TRAF1/2/D1, BIRC2/3 and TAB2 identified by mass spectrometry. TRAF signaling mediates endogenous 4-1BB signal transduction, and the importance of TRAF1^19^ and TRAF2^44^ leading to both canonical and non-canonical NFkB activation^20^ downstream of bbζCAR activation has been established. By QMI, CAR_TRAF2 and multiple TRAF-containing PiSCES appeared to be reduced by CAR stimulation, but IP-mass spec and IP-western blots by us and others^44^ show increases in the amount of CAR_TRAF2. This discrepancy may highlight a known caveat of probing protein complexes in the native state: antibody accessibility may be reduced as signaling complexes form. Collectively, our data support a model that, prior to CAR ligation, CARs are associated with trimers consisting largely of TRAF2. Following CAR activation, the ratio of TRAF1/2 bound to the CAR changes, and while the absolute amount of TRAF2 co-associated with CAR increases (by western blot and mass spec), the accessibility of QMI probe antibodies to bind TRAF2 decreases. Since the QMI probe antibody binds in the TRAF2 Zn-finger domain adjacent to the RING domain, which is the cite of binding-partner recruitment and ubiquitin conjugation following TRAF activation^51,52^, these data are consistent with an activation event. The use of different probes for TRAF2 may confirm this hypothesis, but we have not yet identified suitable antibodies.

Many open questions remain about how CARs are able to engage TRAFs, particularly since natural TRAF ligands are trimeric^18,52^, while activated CARs do not have an obvious trimerization mechanism. However, the recruitment of BIRC2/3 implies clustering of TRAF trimers, which may be driven by CAR clustering or, conversely, may induce clustering and signal enhancement of CARs analogous to TCR clustering. Future work to reconcile the monomeric CAR with the trimeric 4-1BB endogenous receptor structure and the hexagonal model of TRAF signaling^52^ is urgently needed to provide structural insights into the mechanisms of costimulation.

The third major signaling component identified by mass spec was cytoskeletal motility; proteins linked to early endocytic vesicle formation (FLOT1/2), microtubule spindle formation (KIFs, MYO) and T cell migration in response to stimulation (DOCK8) were identified. Activated CARs form non-classical immune synapses^53^, but unlike TCRs, they do not rely on LAT to cluster^54^. Perhaps the CAR’s ability to engage cytoskeletal elements allows it to bypass LAT and form non-classical immune synapses is due to its endogenous cytoskeletal engagement. Alternatively, these data could reflect the sequestration of activated CAR into endosomal compartments for degradation^44^.

Other signaling molecules were also identified by mass spectrometry or QMI that do not fit cleanly into either category. PKCβ (identified by mass spec) and PKCΦ (identified by QMI) are both members of the protein kinase C family, activated by DAG and Ca^2+^. PCKΦ plays a major role in NFkB activation during T cell activation^12^, and PCKβ directly phosphorylates CARD11/CARMA1, promoting the recruitment of the BCL/MALT10 complex and TAK1 to activate NFKB^55^. UBASH3A interacts with CBL-B CD3ζ and ZAP70 in stimulated T cells^24^ and may contribute to CAR endocytosis and degradation following activation. Finally, SHP2, which was already a member of the QMI TCR panel^21^, was found to associate with the CAR, and to reduce its association following stimulation. A complex containing Themis-SHP1 was recently reported to co-associate with bbζCARs, with a requirement for a 10-AA region that overlaps with one of two TRAF2 binding sites^56^, and SHP2 may bind in a similar fashion. While the specific interactors we identified require further investigation, it is striking that the CAR signaling network is reminiscent of a recent human T cell signaling network based on a nonbiased CRISPR screen, which identified both classical TCR signaling molecules and NFkB pathway regulators as critical to T cell signaling^33^.

While we sought to define a bbζCAR signalosome common to all bbζCARs, we observed moderate batch- and CAR-dependent variability. This variability has important clinical implications, as batch-to-batch differences in efficacy and toxicity profiles complicate treatment outcomes. We found that CAR_ZAP70 activation at 5 minutes corelated with the amount of IL2 produced in an overnight assay, which confirms that signalosome activation impacts functional outcomes. More complex clinical outcomes, such as CAR persistence or survival, could potentially also correlate with signalosome measures unique to each batch. For example, differing ratios of TRAF1 to TRAF2, the former of which is only expressed after T cell activation^57^, could affect CAR performance. SNPs in TRAF1 affect the levels of TRAF1 production and contribute to the risk of rheumatic disease^38^, and the levels of TRAF1 can influence activation of canonical vs. non-canonical NFkB activation downstream of 41BB stimulation^58^.

Multiplexed measurement of protein signaling networks in CAR T cells combined with detailed analysis of patient outcomes could allow the identification of protein network features that predict an optimal outcome, and allow the rational design of CAR features or the engineering of intracellular environments that lead to optimal functional performance.

## Materials and Methods

### Cell Culture

CAR T cells and MOCK T cells were prepared using an 8 to 12 day expansion protocol adapted from clinical production practices. Peripheral blood mononuclear cells (PBMCs) were harvested from Leukocyte Reduction System (LRS) Cones provided by STEMCELL Technologies and cryopreserved. Cells were thawed into XVIVO media (Lonza) supplemented with IL-2 at 4.6ng/ml, IL-7 at 5ng/ml, IL-15 at 0.5ng/ml and IL-21 at 1ng/ml (Miltenyi Biotec) at a density of 1 × 10^6^ cells/ml, and activated with Human T-Activator CD3/CD28 Dynabeads™ (ThermoFisher). On Day 1 cells were concentrated to 4 × 10^6^/ml then either transduced with lentiviral vectors encoding CAR constructs at an MOI of 2, with 25ug/ml protamine sulfate (Sigma-Aldrich) or vectorless media with 25ug/ml protamine sulfate. 24 hrs later cells were moved to GREX 24 Well plates (Wilson Wolf) for further expansion at 1 × 10^6^/ml in XVIVO media with cytokines until Dynabeads were removed via magnet on Day 7. Cells were propagated until harvest on day 8-12. Jurkats, CARKats (Jurkats stably transduced with CD19CAR), parental K562s and K562s expressing either CD19 or OKT3 were cultivated in 10% heat-inactivated FBS, 1% Glutaraldehyde, 1% Penicillin-streptomycin, 1% Hepes RPMI 1640 in 175ml non-TC treated flasks.

### Target Cell Fixation

Antigen presenting cells (APCs) for stimulation of primary cells were washed twice in ice-cold PBS then fixed for 30 seconds in 4 ml PBS with 0.1% glutaraldehyde (Sigma) on ice. Immediately followed by quenching with 16 ml 200mM glycine. Cells were washed twice more in ice cold PBS, counted, and mixed with CAR T cells for stimulation.

### Cell stimulation

#### Platebound

CD19 recombinant protein (R&D Systems) or anti-CD3 (Biolegend) with anti-CD28 (Biolgend) antibodies was annealed to wells of 6-well TC coated plates overnight with PBS added to wells as a control. 3×106 CAR or Mock T cells were washed in ice-cold PBS twice then added to plate, followed by a one minute centrifugation at 100g to bring cells in contact with plate bound protein. Plate placed in 37C water bath for 5 minutes followed by immediate lysis in 1% digitonin lysis buffer on ice.

#### K562 Targets

CAR T and target cells (fixed or unfixed) were washed in ice-cold PBS twice, then combined at a 1:1 ratio and mixed. Cells were spun at 300g × 5 min at 4C, the PBS removed, and pellet was agitated by wash-boarding the tube across an uneven surface. Cell pellets were stimulated by immersing tubes in a 37C waterbath for 5 minutes, followed freezing in liquid N_2_. Cell pellets were stored at −80 until further processing.

### Lysis

Frozen cell pellets were resuspended in 250-500ul lysis buffer (1% either Digitonin, 1x phosphatase inhibitor cocktail (Sigma), 1x protease inhibitor cocktail (Sigma), 1x sodium orthovanadate (Sigma) and 1x sodium fluorine (Sigma) in 50mm TRIS, 150mm NaCl, pH=7.4). Lysate was incubated on ice for 15 minutes and spun down at 13000g for 10 min at 4 C. Supernatant was harvested and used for IP of protein targets.

### Co-immunoprecipitation for mass spectrometry

Cryopreserved CAR T cells produced from heathy donors in the Seattle Children’s GMP production facility were thawed and cultured overnight in XVIVO media (Lonza) supplemented with IL-2 at 4.6ng/ml, IL-7 at 5ng/ml, IL-15 at 0.5ng/ml and IL-21 at 1ng/ml (Miltenyi Biotec). The following day, cells were mixed 1:1 with fixed target cells (as above) and either warmed for 5 minutes to 37C to allow signal transduction, or maintained on ice to control for scFv-CD19 binding without signal transduction. Cells were lysed 1%NP40 lysis buffer. Protein G magnetic beads (NEB) were incubated with anti-2A (Novus) for 40 minutes with agitation at RT, followed by cross linking of antibody to beads by incubation in 25mM DMP for 45 minutes with agitation at RT. Cross-linking reaction quenched with 50mM Tris-HCl. Cell lysates were incubated overnight with 2A-proteinG beads, washed twice in lysis buffer, twice in lysis buffer without detergent, and proteins were eluted in 200mM glycine pH 2. Eluted proteins were precipitated using methanol-chloroform and pellets were sent to mass spectrometry for processing.

### Co-Immunoprecipitation for Western Blot

Protein G beads covalently bound and cross-linked to 2A antibodies for CO-IPs to be ran on western blot were prepared following MS sample preparation with the addition of a final elution of excess un-cross-linked antibody with 200mM glycine pH 2. Eluted proteins were precipitated in acetone. Additionally, immunoprecipitation for westerns was performed on cell lysates derived from APCs mixed with CAR and Mock Transduced primary T cells from LRS Cones (STEMCELL).

### Mass Spectrometry

Protein pellets were resuspended in 20 µL of 8 M urea in 100 mM ammonium bicarbonate and vortexed thoroughly. Protein disulfide bonds were reduced by the addition of tris (2-carboxyethyl) phosphine (TCEP) to a final concentration of 20 mM and incubated at room temperature for 20 min. Cysteines were alkylated by the addition of 2-chloroacetamide to a final concentration of 20 mM and incubating at room temperature for 30 min. Proteolytic digestion was initiated by the addition of 250 ng of endoproteinase Lys-C (Promega) and incubating at room temperature for 2 hours. The proteolytic digestion was continued by the addition of 500 ng of trypsin (Promega) and incubating overnight at 37°C with gentle shaking. Digestions were stopped by the addition of trifluoroacetic acid (TFA) to a final concentration of 0.1%. The resulting peptide samples were desalted on a C18 Ultra-micro Spin Column (Harvard Apparatus) using 70% acetonitrile in 0.1% TFA and taken to dryness by vacuum centrifugation.

Dried samples were brought up in 20 µL of 2% acetonitrile in 0.1% formic acid and 5 µL was analyzed by LC/ESI MS/MS with a ThermoFisher Scientific Easy1000 nLC coupled to an Orbitrap Fusion mass spectrometer. In-line de-salting was accomplished using a reversed-phase trap column (100 μm × 20 mm) packed with Magic C_18_AQ (5-μm, 200Å resin; Michrom Bioresources) followed by peptide separations on a reversed-phase column (75 μm × 270 mm) packed with ReproSil-Pur C_18_AQ (3-μm, 120Å resin; Dr. Maisch) directly mounted on the electrospray ion source. Chromatographic separation was carried out adjusting the elution gradient from 5% to 28% B (80% acetonitrile with 20% water and 0.1% formic acid) in 90 minutes, 28% to 50% B in 10 min, holding at 50% B for 3 min, 50% to 95% B in 2 min, and holding at 95% B for 1 minute. A flow rate of 300 nL/minute was used for chromatographic separations with the temperature of the chromatographic column maintained at 40 ℃. A spray voltage of 2200 V was applied to the electrospray tip while the Orbitrap Eclipse instrument was operated in the data-dependent mode. MS survey scans were in the Orbitrap (AGC target value of 5E5, resolution 120,000, and max injection time 50 ms) using a scan range of 400 m/z to 1500 m/z and a 3 sec cycle time. MS/MS spectra were acquired in the linear ion trap (AGC target value of 1E4, rapid scan rate, and max injection time 45 ms) with an isolation window of 1.6 and using higher energy collision-induced dissociation (HCD) activation with a collision energy of 27%. The dynamic exclusion duration was set to 20 sec.

Protein database searching and lablel-free quantification (LFQ) were performed using ThermoFisher Scientific Proteome Discoverer v2.4. The data were searched against a Uniprot human database (UP000005640; downloaded 120119) that included common contaminants (cRAP; Global Proteome Machine). Search setting included the proteolytic enzyme set to trypsin, maximum missed cleavages set to 2, precursor ion tolerance set to 10 ppm, and the fragment ion tolerance set to 0.6 Da. Dynamic modifications were set to oxidation on methionine (+15.995 Da), phosphorylation on serine and threonine (+79.966), acetylation of the protein N-terminus (+42.011 Da), methionine loss at the protein N-terminus (−131.040 Da), and methionine loss and acetylation of the protein N-terminuns (−89.030 Da). A static modification of carbamidomethylation of cysteine (+57.021 Da) was used. Sequest HT was used for protein database searching and Percolator was used for peptide validation. Peptide to spectrum matches (PSMs) were filtered to a 1% false discovery rate and the resulting proteins were further filtered to a 1% false discovery rate. LFQ analysis was carried out using the Precursor Ion Quantifier node using the default settings. Normalization was carried out using the total peptide amount setting.

### Screening QMI antibodies with IP-FCM

Performed as described previously^59^. CML Beads (CML Latex Microspheres, Invitrogen, USA) were activated with EDAC (1-ethyl-3-(3-dimethylaminopropyl) carbodiimide HCL) then coupled to 50ul of antibody at 0.5 mg/ml for 2 hours at room temperature with agitation at 1400rpm. Probe antibodies were biotinylated at 50-fold molar excess with EZ-link Sulfo-NHS-Biotin (Thermo, USA). Coupled CML beads were added to lysate of frozen cell pellets at equal bead to protein quantity ratios across conditions then incubated at 4°C overnight with rotation. Beads and any bound protein complexes were then washed three times in Fly-P buffer before being distributed across a 96 well plate, two times for every biotinylated probe antibody to be used. Probes were then added at 2.5ug/ml and the plate was incubated with agitation at 600rpm for 1hr at 4°C. Two washed in Fly-P buffer followed by incubation in 10ug/ml streptavidin-phycoerythrin (PE, Biolegend) for 30 minutes. After three washes in Fly-P, CML beads were analyzed for PE fluorescence on a flow cytometer (Novocyte). Data consisted of MFI and bead distributions.

### QMI

QMI experiments were performed as described previously^21,32^. All experiments and procedures were performed at 4°C or on ice, starting with the preparation of a master mix of an equal number of each antibody-coupled Luminex bead used for IPs. Equal amounts of bead master mix was distributed into cell lysates whose protein concentrations were normalized by BCA assay. IPs of protein complexes were performed overnight with rotation then washed twice in Fly-P buffer and distributed across a 96 well plate at two wells per detection antibody. Biotinylated or PE-conjugated detection antibodies diluted to 2.5ug/ml were added to the plate for 1 hr with shaking at 600 rpm. See table S2 for IP and detection antibody details. After three washes in Fly-P buffer using the Bio-Plex Pro II magnetic plate washer, microbeads and captured protein complexes stained with detection antibody were incubated in streptavidin-PE or Fly-P buffer for 30 minutes, then washed again as before and resuspended in 120ul Fly-P buffer. Fluorescence was analyzed and data acquired via a customized, refrigerated Bio-Plex 200 using Bio-plex Manager software (v.6.2). XML formatted data files exported for further analysis.

### QMI Data Analysis

A detailed video protocol and source code for QMI data analysis has been previously published^32^, including statistical code run in MatLab (ANC) or R (CNA). Briefly, for CNA, data were normalized using the ComBAT function, PiSCES were organized into modules based on correlated behavior across experimental conditions and replicates, and modules were then correlated with experimental variables. PiSCES that were significantly (P<0.05) and strongly (module membership < 0.7) correlated to a module that was itself significantly correlated to an experimental variable were considered “CNA-significant”. Modules that correlated with CD3 stimulation and contained PI3K_LAT were re-named “turquoise”, and modules that correlated with CAR expression and contained CAR-containing interactions were re-named “black”, to highlight the fact that while the specific interactions that comprised each module varied by experiment, the same overall patterns were observed between experiments, similar to previous reports in RNA expression^60^. Meanwhile, for ANC, individual PiSCES were compared for each experimental N between the stimulated group and the control group using nonparametric Kolomogrov-Schmirnov statistics corrected to maintain a type 1 error rate of 0.05, corrected for multiple comparisons using Bonferroni correction. For equations, see ^21^. Only PiSCES that were significant by both ANC and CNA are displayed in heatmaps and node-edge diagrams, generated in R and cytoscape, respectively.

### Western blot

All western blot experiments consisted of proteins denatured in SDS sample buffer (4x, Bio-rad) with 10% (v/v) beta-mercaptoethanol and loaded in 10% acrylamide gels. Gels were transferred onto polyvinylidene difluoride (Millipore) membranes then blocked in 4% milk TBST (0.05 M Tris, 0.15 M NaCl, pH7.2, 0.1% (v/v) Tween20) for 60 min at room temperature with gentle rocking. Primary antibodies applied overnight at 4°C in 4% milk. The following primary antibodies were used, all at a 1:1000 volumetric dilution: anti-2A (3H4, Novus), anti-TRAF2 (F-2, Santa Cruz), anti-ZAP70 (D9H10, Novus), and TAK1 (28H25L68, Thermo). Blots were probed with species specific horseradish peroxidase conjugated antibodies then imaged using SuperSignal West Femto (thermofisher) substrate on a Protein Simple imaging system.

### IL-2 ELISA Cytokine Release Assay

Effector and target cells were combined at various ratios in XVIVO media without cytokines at 1 × 10^6^ effector cells/ml and incubated overnight at 37 C, 5% CO_2_ in 96-well non-TC plates. Cells were then spun down at 300g for five min and supernatant frozen at −80 C. IL-2 concentration in harvested supernatant was then analyzed using Human IL-2 ELISA Max Kit (BioLegend) per provided instruction. All cell effector activations were performed in four separate replicates and analyzed as separate samples. A SpectraMax i3x plate reader was used for spectrometry and data generation.

## Acknowledgements

We thank Brendan K Reed, Juliane Gust, and the entire SEPS lab for helpful discussions and technical assistance. We gratefully acknowledge the following individuals for providing cell lines or reagents: Scott Cramer, Jessica L. Maiers, Christine Mayr, Harald Wajant, Tania H Watts and Nicolas Bidère. We also thank Meredith A. Jackson for providing protocols and CAR constructs.

## Supplementary Material

**Table S1:**
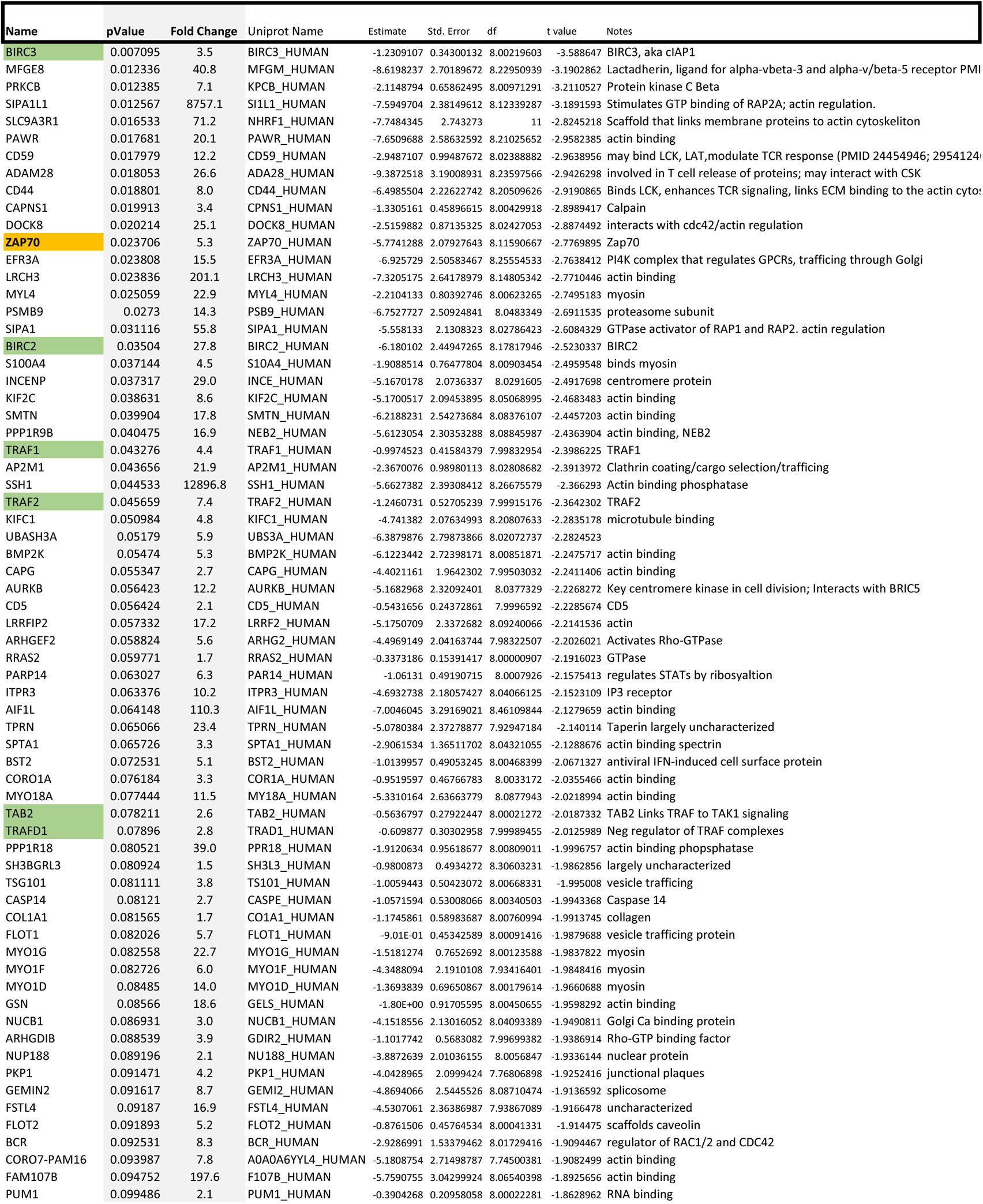
Protein identified by IP-mass spectrometry as significantly enriched in stimulated vs. unstimulated bbζCAR immunoprecipitates.

**Figure S1:**
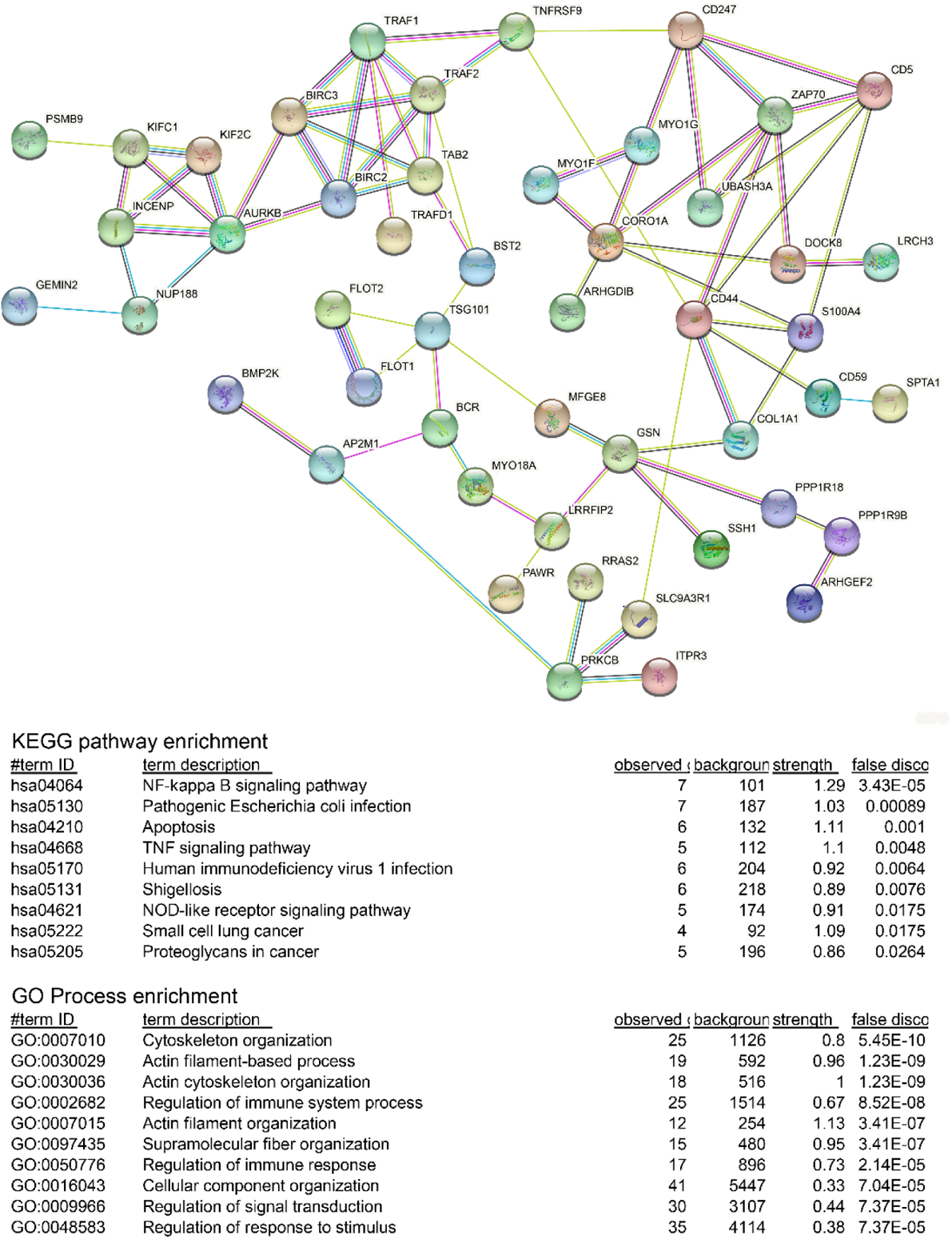
Known interactions in the STRING database among proteins identified by LC/MS/MS of bbζCAR immunoprecipitates. Proteins were entered into the Sting database with the following settings: Full STRING network, all interaction sources, medium confidence. KEGG pathway enrichment and GO process enrichment are shown as generated by STRING.

**Figure S2:**
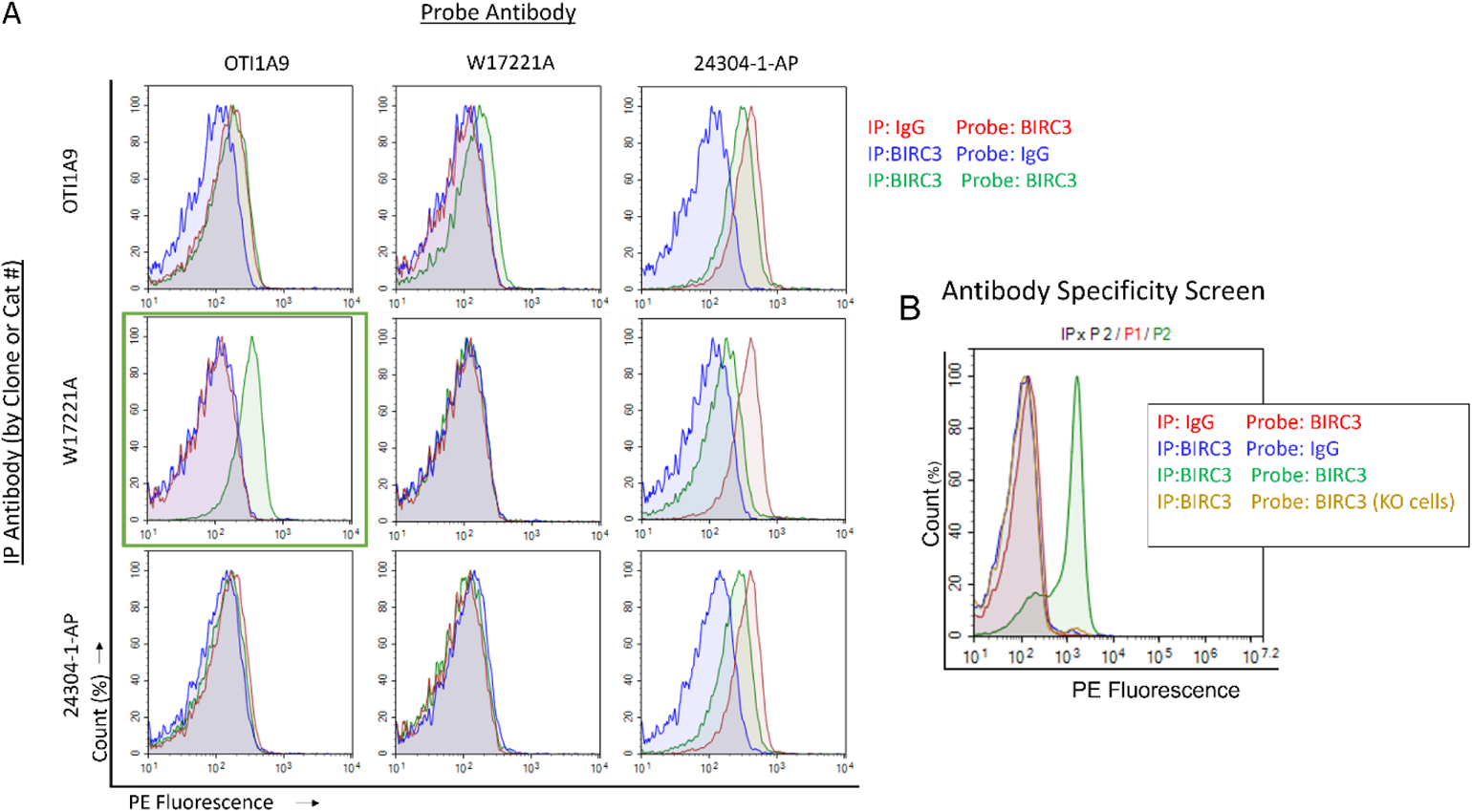
Identification of compatible antibodies for QMI. A) Antibodies were coupled to 5um CML beads, or biotinylated for use as probes. Coupled beads were mixed with Jurkat cell lysate, washed, and incubated with each probe antibody. IP_Probe combinations that produced the strongest fluorescent signal (green histograms) over IgG IP and probe controls (red and blue histograms, respectively) were selected (green box). B) The selected pairs were validated using knockout cell lysate-here RAJI cells with BIRC3 knocked out by CRISPR^1^. If median fluorescent intensity was reduced by >90%, the pair was considered ‘validated’. The antibodies shown are selected examples; for the full list of IPs, Probes and validation experiments, see Table S2.

**Table S2:**
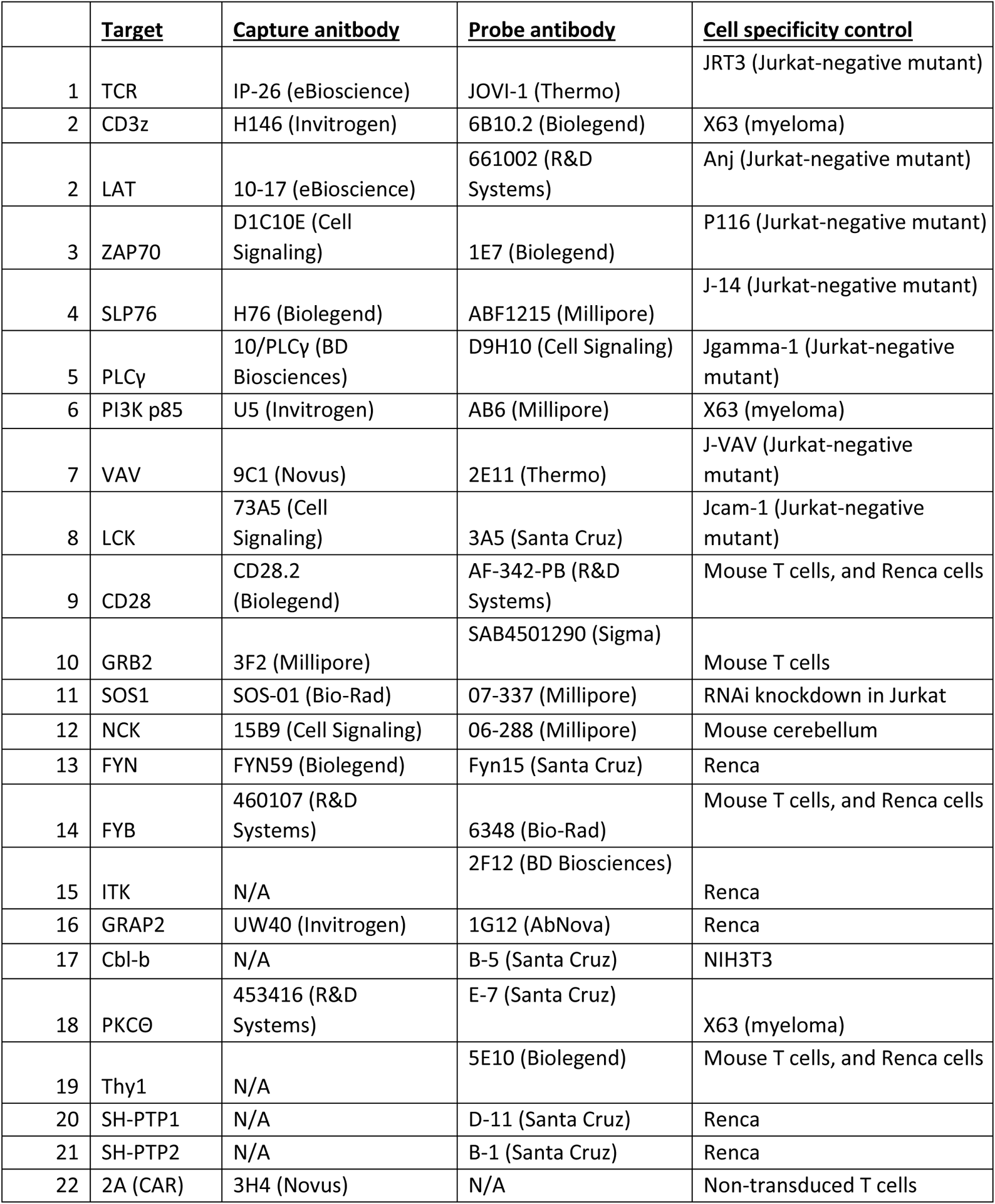
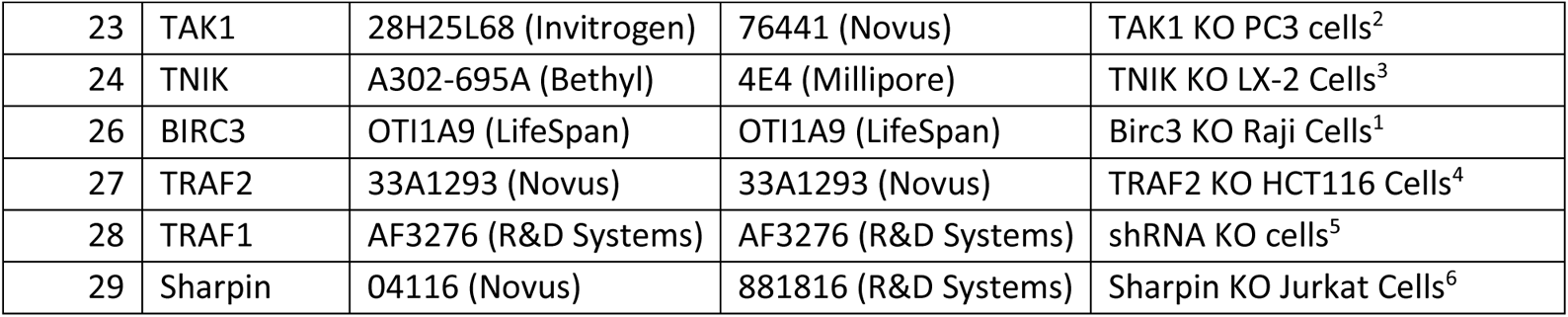
IP and probe antibodies used in QMI experiments, and the material used for validation studies. See Figure S2.

**Figure S3:**
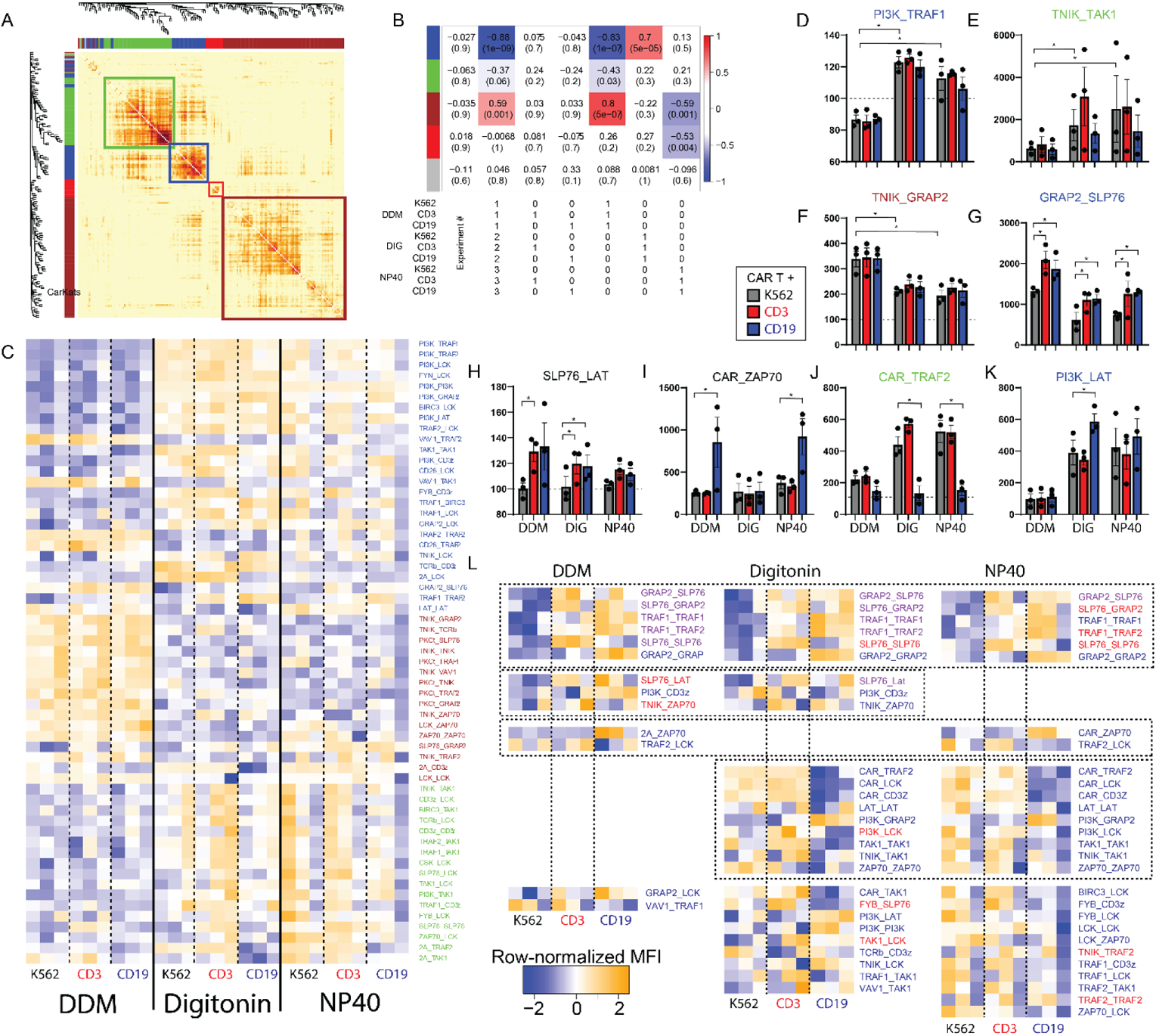
Detergent optimization of bbζCAR QMI. Jurkat cells stably expressing CD19 bbζCAR were mixed 1:1 with K562 cells expressing nothing, anti-CD3 or anti-CD19 and lysed in DDM, Digitonin, or NP40, and QMI was performed. A) Correlation network analysis clustered all detected PiSCES by their correlated behavior across N=27 individual measurements, and identified color-coded modules of correlated interactions. B) A module-trait correlation table reports the correlation coefficient and p-value (in parenthesis) between the eigenvector of each color-coded CNA module (colored boxes) and a binary description of experimental variables, below table. Module-trait correlations greater than 0.4 are highlighted with red (positive correlation) or blue (negative correlation). C) Row-normalized heatmap of ANCՈCNA-significant proteins in shared complexes (PiSCES) in the Blue, Brown or Green modules. D) The blue module, anti-correlated with DDM, contained PiSCES such as PI3K_TRAF1, which were detected in DIG and NP40 but poorly detected in DDM. * p<0.05 by ANC. E) The green module, exemplified by TNIK_TAK1, was similar, but showed higher inter-experimental variability. F) The brown module, exemplified by TNIK_GRAP2, was most strongly detected in DDM. G) GRAP2_SLP76 was increased following CD3 or CD19 stimulation in all detergents. H) SLP76_LAT was increased following CD3 or CD19 stimulation in DDM o DIG, but not NP40. I) CAR_ZAP70 was increased with CD19 stimulation in DDM or NP40, but not DIG. J) CAR_TRAF2 was decreased with CD19 stimulation in DIG or NP40, but not detected in DDM. K) PI3K_LAT was increased by CAR stimulation only in DIG. L) Row-normalized heatmap comparing PiSCES that were ANC-significant for a comparison between CAR-K562 and CAR-CD3 (red) or CAR-CD19 (blue) or both (purple) in each detergent. The rows are arranged by PiSCES that were significant in all three detergents (top), or 2/3 (middle), or only a single comparison (bottom), as indicated by dashed boxes.

**Figure S4:**
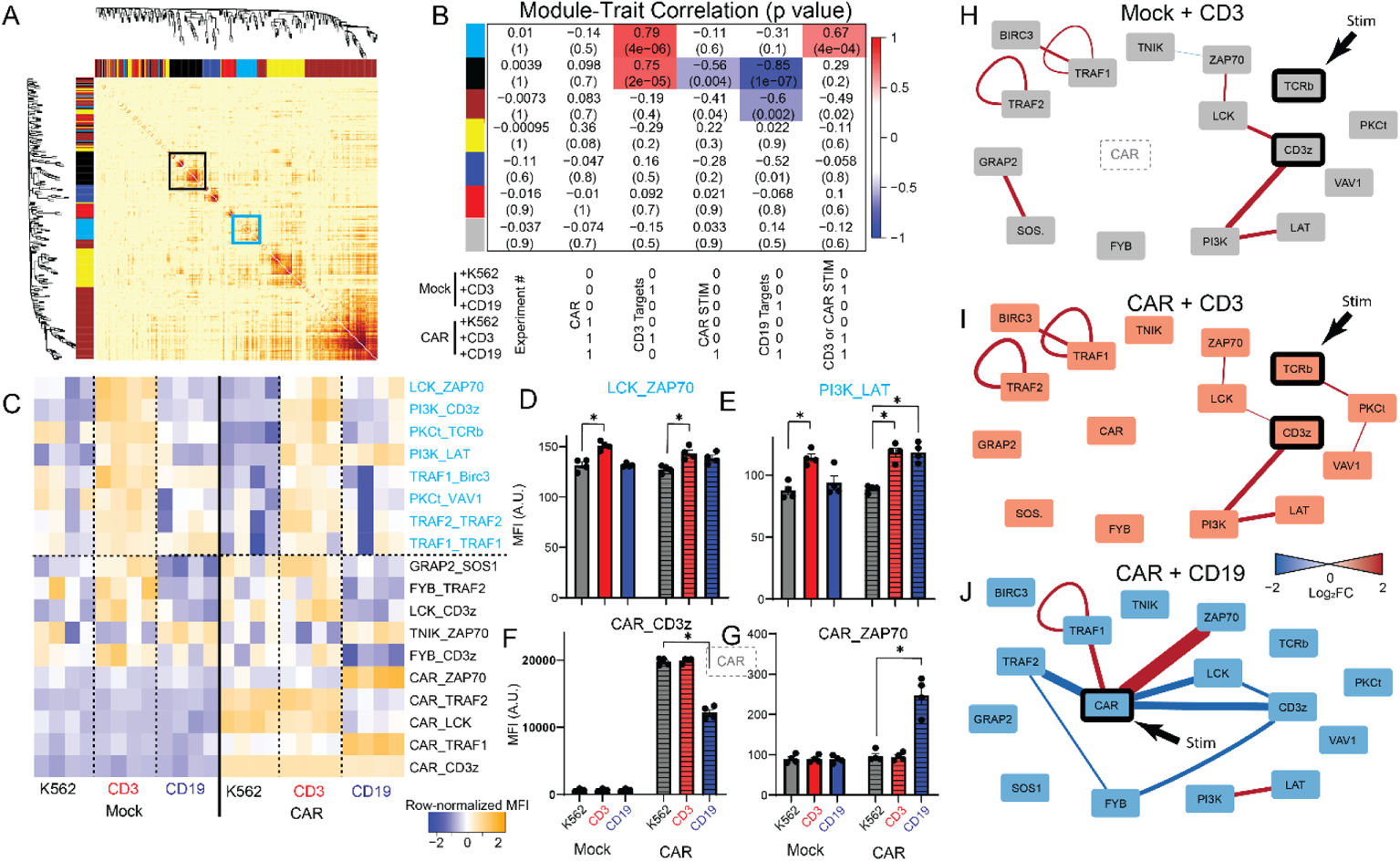
bbζCAR activates TCR- and TRAF-associated signaling networks when target cells are not fixed. Related to Figure 3. bbζCAR or mock-transduced T cells were stimulated with *<u>live</u>* K562 targets expressing nothing, anti-CD3 (OKT3) or CD19 for 5 minutes. A) Correlation network analysis clustered all detected PiSCES by their correlated behavior across N=24 individual measurements, and identified color-coded modules of correlated interactions. B) A module-trait correlation table reports the correlation coefficient and p-value (in parenthesis) between the eigenvector of each color-coded CNA module (colored boxes) and a binary description of experimental variables, below table. Module-trait correlations greater than 0.6 are highlighted with red (positive correlation) or blue (negative correlation). C) Row-normalized heatmap of ANCՈCNA-significant proteins in shared complexes (PiSCES) in the turquoise or black modules. D-G) MFI of (D) LCK_ZAP70 and (E) PI3K_LAT representative of the Turquoise module, which increases following anti-CD3 stimulation and increases to a lesser degree following CD19 stimulation. * p<0.05 by ANC. MFI of (F) CAR_CD3 and (G) CAR_ZAP70, both in the Black module. H-J) PiSCES that changed with anti-CD3 stimulation in H) mock-transduced or I) CAR t cells, or that changed with J) CD19 stimulation in CAR T cells. Edges represent a change in the amount of interaction between the connected nodes; edge color and width indicate direction and magnitude of the change.

**Figure S5:**
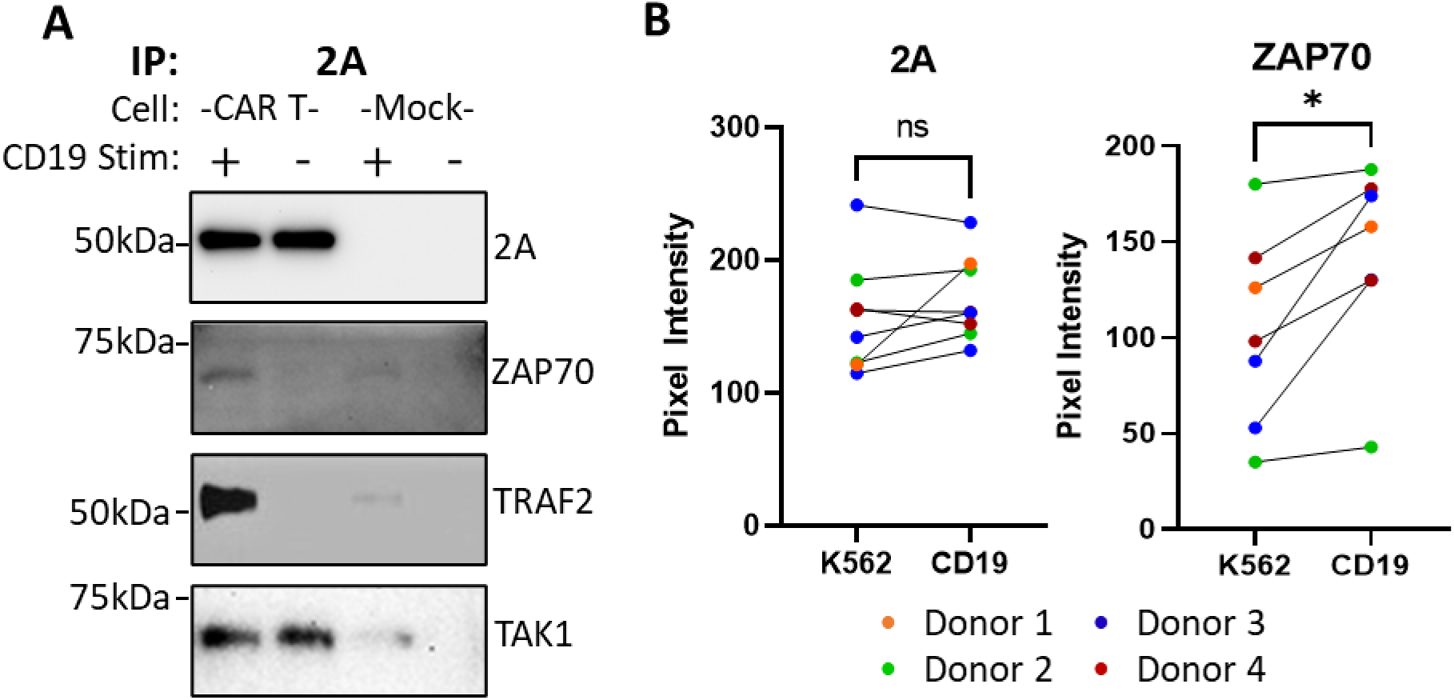
Co-immunoprecipitations of bbζCAR with ZAP70, TRAF2 and TAK1. A) The bbζCAR was IP’d via the 2A scar from unstimulated (K562) and stimulated (CD19) CAR T cells, and blots were probed with 2A, ZAP70, TRAF2 or TAK1. Blots shown represent N=2-8 separate experiments. B) Pixel intensity of 2A and ZAP70 western blot bands compared across multiple batches of CAR T cells from four donors, showing significantly increased ZAP70 recruitment with stimulation.

## References

1. Lundby, A. et al. Oncogenic Mutations Rewire Signaling Pathways by Switching Protein Recruitment to Phosphotyrosine Sites. Cell 179, 543–560.e26 (2019).

2. Wu, L., Wei, Q., Brzostek, J. & Gascoigne, N. R. J. Signaling from T cell receptors (TCRs) and chimeric antigen receptors (CARs) on T cells. Cell. Mol. Immunol. 17, 600–612 (2020).

3. Sadelain, M., Rivière, I. & Riddell, S. Therapeutic T cell engineering. Nature 545, 423–431 (2017).

4. Maude, S. L. et al. Tisagenlecleucel in Children and Young Adults with B-Cell Lymphoblastic Leukemia. N. Engl. J. Med. 378, 439–448 (2018).

5. Jaklevic, M. C. CAR-T Therapy Is Approved for Non-Hodgkin Lymphoma. JAMA 325, 1032 (2021).

6. Neelapu, S. S. et al. Axicabtagene Ciloleucel CAR T-Cell Therapy in Refractory Large B-Cell Lymphoma. N. Engl. J. Med. 377, 2531–2544 (2017).

7. Frey, N. V. Approval of brexucabtagene autoleucel for adults with relapsed and refractory acute lymphocytic leukemia. Blood (2022) doi:10.1182/blood.2021014892.

8. Gardner, R. A. et al. Intent-to-treat leukemia remission by CD19 CAR T cells of defined formulation and dose in children and young adults. Blood 129, 3322–3331 (2017).

9. Martinez, M. & Moon, E. K. CAR T Cells for Solid Tumors: New Strategies for Finding, Infiltrating, and Surviving in the Tumor Microenvironment. Front. Immunol. 10, 128 (2019).

10. Gust, J., Ponce, R., Liles, W. C., Garden, G. A. & Turtle, C. J. Cytokines in CAR T Cell-Associated Neurotoxicity. Front. Immunol. 11, 577027 (2020).

11. Schrum, A. G. & Gil, D. Robustness and Specificity in Signal Transduction via Physiologic Protein Interaction Networks. Clin. Exp. Pharmacol. 2, S3.001 (2012).

12. Shah, K., Al-Haidari, A., Sun, J. & Kazi, J. U. T cell receptor (TCR) signaling in health and disease. Signal Transduct. Target. Ther. 6, 1–26 (2021).

13. de la Cruz, J. et al. Basal and antigen-induced exposure of the proline-rich sequence in CD3ε. J. Immunol. Baltim. Md 1950 186, 2282–2290 (2011).

14. Neier, S. C. et al. The early proximal αβ TCR signalosome specifies thymic selection outcome through a quantitative protein interaction network. Sci. Immunol. 4, (2019).

15. Gudipati, V. et al. Inefficient CAR-proximal signaling blunts antigen sensitivity. Nat. Immunol. 21, 848–856 (2020).

16. Salter, A. I. et al. Phosphoproteomic analysis of chimeric antigen receptor signaling reveals kinetic and quantitative differences that affect cell function. Sci. Signal. 11, eaat6753 (2018).

17. Salter, A. I. et al. Comparative analysis of TCR and CAR signaling informs CAR designs with superior antigen sensitivity and in vivo function. Sci. Signal. 14, eabe2606 (2021).

18. Zapata, J. M. et al. CD137 (4-1BB) Signalosome: Complexity Is a Matter of TRAFs. Front. Immunol. 9, (2018).

19. Li, G. et al. 4-1BB enhancement of CAR T function requires NF-κB and TRAFs. JCI Insight 3, 121322 (2018).

20. Philipson, B. I. et al. 4-1BB costimulation promotes CAR T cell survival through noncanonical NF-κB signaling. Sci. Signal. 13, eaay8248 (2020).

21. Smith, S. E. P. et al. Multiplex matrix network analysis of protein complexes in the human TCR signalosome. Sci. Signal. 9, rs7 (2016).

22. Mellacheruvu, D. et al. The CRAPome: a contaminant repository for affinity purification-mass spectrometry data. Nat. Methods 10, 730–736 (2013).

23. Shinohara, H. et al. PKC beta regulates BCR-mediated IKK activation by facilitating the interaction between TAK1 and CARMA1. J. Exp. Med. 202, 1423–1431 (2005).

24. Ge, Y., Paisie, T. K., Chen, S. & Concannon, P. UBASH3A Regulates the Synthesis and Dynamics of TCR–CD3 Complexes. J. Immunol. 203, 2827–2836 (2019).

25. Matson, C. A. et al. CD5 dynamically calibrates basal NF-κB signaling in T cells during thymic development and peripheral activation. Proc. Natl. Acad. Sci. 117, 14342–14353 (2020).

26. Voisinne, G., Gonzalez de Peredo, A. & Roncagalli, R. CD5, an Undercover Regulator of TCR Signaling. Front. Immunol. 9, (2018).

27. Föger, N., Marhaba, R. & Zöller, M. CD44 supports T cell proliferation and apoptosis by apposition of protein kinases. Eur. J. Immunol. 30, 2888–2899 (2000).

28. Dimitrakopoulos, G. N., Klapa, M. I. & Moschonas, N. K. PICKLE 3.0: Enriching the human Meta-database with the mouse protein interactome extended via mouse-human orthology. Bioinforma. Oxf. Engl. btaa1070 (2020) doi:10.1093/bioinformatics/btaa1070.

29. Szklarczyk, D. et al. The STRING database in 2021: customizable protein-protein networks, and functional characterization of user-uploaded gene/measurement sets. Nucleic Acids Res. 49, D605–D612 (2021).

30. Xu, X. et al. LRCH1 interferes with DOCK8-Cdc42-induced T cell migration and ameliorates experimental autoimmune encephalomyelitis. J. Exp. Med. 214, 209–226 (2017).

31. Ham, H., Huynh, W., Schoon, R. A., Vale, R. D. & Billadeau, D. D. HkRP3 is a microtubule-binding protein regulating lytic granule clustering and NK cell killing. J. Immunol. Baltim. Md 1950 194, 3984–3996 (2015).

32. Brown, E. A., Neier, S. C., Neuhauser, C., Schrum, A. G. & Smith, S. E. P. Quantification of Protein Interaction Network Dynamics using Multiplexed Co-Immunoprecipitation. JoVE J. Vis. Exp. e60029 (2019) doi:10.3791/60029.

33. Schmidt, R. et al. CRISPR activation and interference screens decode stimulation responses in primary human T cells. Science 375, eabj4008 (2022).

34. Jaeger-Ruckstuhl, C. A. et al. TNIK signaling imprints CD8+ T cell memory formation early after priming. Nat. Commun. 11, 1632 (2020).

35. Ikeda, F. et al. SHARPIN forms a linear ubiquitin ligase complex regulating NF-κB activity and apoptosis. Nature 471, 637–641 (2011).

36. Tokunaga, F. et al. SHARPIN is a component of the NF-κB-activating linear ubiquitin chain assembly complex. Nature 471, 633–636 (2011).

37. Oikawa, D., Hatanaka, N., Suzuki, T. & Tokunaga, F. Cellular and Mathematical Analyses of LUBAC Involvement in T Cell Receptor-Mediated NF-κB Activation Pathway. Front. Immunol. 11, 601926 (2020).

38. Abdul-Sater, A. A. et al. The signaling adaptor TRAF1 negatively regulates Toll-like receptor signaling and this underlies its role in rheumatic disease. Nat. Immunol. 18, 26–35 (2017).

39. Lautz, J. D., Brown, E. A., Williams VanSchoiack, A. A. & Smith, S. E. P. Synaptic activity induces input-specific rearrangements in a targeted synaptic protein interaction network. J. Neurochem. 146, 540–559 (2018).

40. Lautz, J. D. et al. Activity-dependent changes in synaptic protein complex composition are consistent in different detergents despite differential solubility. Sci. Rep. 9, 10890 (2019).

41. Langfelder, P. & Horvath, S. WGCNA: an R package for weighted correlation network analysis. BMC Bioinformatics 9, 559 (2008).

42. van Oers, N. S., von Boehmer, H. & Weiss, A. The pre-T cell receptor (TCR) complex is functionally coupled to the TCR-zeta subunit. J. Exp. Med. 182, 1585–1590 (1995).

43. Ramello, M. C. et al. An immunoproteomic approach to characterize the CAR interactome and signalosome. Sci. Signal. 12, eaap9777 (2019).

44. Li, W. et al. Chimeric Antigen Receptor Designed to Prevent Ubiquitination and Downregulation Showed Durable Antitumor Efficacy. Immunity 53, 456–470.e6 (2020).

45. Singh, N. et al. Antigen-independent activation enhances the efficacy of 4-1BB-costimulated CD22 CAR T cells. Nat. Med. 27, 842–850 (2021).

46. Levine, B. L., Miskin, J., Wonnacott, K. & Keir, C. Global Manufacturing of CAR T Cell Therapy. Mol. Ther. Methods Clin. Dev. 4, 92–101 (2016).

47. Ravanpay, A. C. et al. EGFR806-CAR T cells selectively target a tumor-restricted EGFR epitope in glioblastoma. Oncotarget 10, 7080–7095 (2019).

48. Wei, Q. et al. Lck bound to coreceptor is less active than free Lck. Proc. Natl. Acad. Sci. 117, 15809–15817 (2020).

49. Li, L. et al. Ionic CD3−Lck interaction regulates the initiation of T-cell receptor signaling. Proc. Natl. Acad. Sci. 114, E5891–E5899 (2017).

50. Lo, W.-L. et al. Lck promotes Zap70-dependent LAT phosphorylation by bridging Zap70 to LAT. Nat. Immunol. 19, 733–741 (2018).

51. Park, H. H. Structure of TRAF Family: Current Understanding of Receptor Recognition. Front. Immunol. 9, (2018).

52. Vanamee, É. S. & Faustman, D. L. Structural principles of tumor necrosis factor superfamily signaling. Sci. Signal. 11, eaao4910 (2018).

53. Davenport, A. J. et al. Chimeric antigen receptor T cells form nonclassical and potent immune synapses driving rapid cytotoxicity. Proc. Natl. Acad. Sci. 115, E2068–E2076 (2018).

54. Dong, R. et al. Rewired signaling network in T cells expressing the chimeric antigen receptor (CAR). EMBO J. 39, e104730 (2020).

55. Kang, S. W. et al. PKCbeta modulates antigen receptor signaling via regulation of Btk membrane localization. EMBO J. 20, 5692–5702 (2001).

56. Sun, C. et al. THEMIS-SHP1 Recruitment by 4-1BB Tunes LCK-Mediated Priming of Chimeric Antigen Receptor-Redirected T Cells. Cancer Cell 37, 216–225.e6 (2020).

57. Edilova, M. I., Abdul-Sater, A. A. & Watts, T. H. TRAF1 Signaling in Human Health and Disease. Front. Immunol. 9, 2969 (2018).

58. McPherson, A. J., Snell, L. M., Mak, T. W. & Watts, T. H. Opposing roles for TRAF1 in the alternative versus classical NF-κB pathway in T cells. J. Biol. Chem. 287, 23010–23019 (2012).

59. Davis, T. R. & Schrum, A. G. IP-FCM: Immunoprecipitation Detected by Flow Cytometry. JoVE J. Vis. Exp. e2066 (2010) doi:10.3791/2066.

60. Voineagu, I. et al. Transcriptomic analysis of autistic brain reveals convergent molecular pathology. Nature 474, 380–384 (2011).

## References

1. Lee, S.-H. & Mayr, C. Gain of Additional BIRC3 Protein Functions through 3ʹ-UTR-Mediated Protein Complex Formation. Mol. Cell 74, 701–712.e9 (2019).

2. Bayne, R. S. et al. MAP3K7 and CHD1 Are Novel Mediators of Resistance to Oncolytic Vesicular Stomatitis Virus in Prostate Cancer Cells. Mol. Ther. - Oncolytics 17, 496–507 (2020).

3. Buchl, S. C. et al. Traf2 and NCK Interacting Kinase Is a Critical Regulator of Procollagen I Trafficking and Hepatic Fibrogenesis in Mice. Hepatol. Commun. 6, 593–609 (2022).

4. Kreckel, J., Anany, M. A., Siegmund, D. & Wajant, H. TRAF2 Controls Death Receptor-Induced Caspase-8 Processing and Facilitates Proinflammatory Signaling. Front. Immunol. 10, (2019).

5. Abdul-Sater, A. A. et al. The signaling adaptor TRAF1 negatively regulates Toll-like receptor signaling and this underlies its role in rheumatic disease. Nat. Immunol. 18, 26–35 (2017).

6. Thys, A. et al. Serine 165 phosphorylation of SHARPIN regulates the activation of NF-κB. iScience 24, 101939 (2021).

